# Fusion pore dynamics of large secretory vesicles define a distinct mechanism of exocytosis

**DOI:** 10.1101/2023.06.08.544180

**Authors:** Tom Biton, Nadav Scher, Shari Carmon, Yael Elbaz-Alon, Eyal D. Schejter, Ben-Zion Shilo, Ori Avinoam

## Abstract

Exocrine cells utilize large secretory vesicles (LSVs) up to 10 μm in diameter. LSVs fuse with the apical surface, often recruiting actomyosin to extrude their content through dynamic fusion pores. The molecular mechanism regulating pore dynamics remains largely uncharacterized. We observe that the fusion pore of LSVs in the Drosophila larval salivary glands expand, stabilize, and constrict. Arp2/3 is essential for pore expansion and stabilization, while myosin II is essential for pore constriction. We identify several Bin-Amphiphysin-Rvs (BAR) homology domain proteins that regulate fusion pore expansion and stabilization. We show that the I-BAR protein Missing-in-Metastasis (MIM) localizes to the fusion site and is essential for pore expansion and stabilization. The MIM I-BAR domain is essential but not sufficient for localization and function. We conclude that MIM acts in concert with actin, myosin II, and additional BAR-domain proteins to control fusion pore dynamics, mediating a distinct mode of exocytosis that facilitates actomyosin-dependent content release that maintains apical membrane homeostasis during secretion.

## Introduction

The fusion of secretory vesicles with the plasma membrane is essential for the exocytotic release of bioactive materials such as neurotransmitters, hormones, and digestive enzymes from neuronal, endocrine, and exocrine cells. The latter, customarily utilize large secretory vesicles (LSVs) with diameters that are 1-2 orders of magnitude larger than those of neuronal or endocrine vesicles, challenging the conventional mechanisms of vesicle exocytosis and recycling.

Secretory vesicles dock and fuse with the cell surface, connecting the vesicle lumen and the extracellular space via a fusion pore (Breckenridge and Almers, 1987; Curran et al., 1993; Hastoy et al., 2017; Sharma and Lindau, 2018) The formation of a fusion pore at the contact site between the vesicle and cell membrane is mediated by soluble N-ethylmaleimide-sensitive fusion attachment protein receptors (SNAREs) (Söllner et al., 1993a; b; Sudhof and Rothman, 2009; Fang et al., 2008; Ngatchou et al., 2010; Wiederhold et al., 2010). Fusion pores are initially nanometric and can flicker open and reseal spontaneously several times before stabilizing, expanding, or constricting (Chanturiya et al., 1997; Curran et al., 1993; Vardjan et al., 2013; Toledo et al., 2018). If the fusion pore reseals, the vesicle is recycled *en bloc*, retaining some of its cargo (i.e. kiss-and-run(Toledo et al., 1993; Rizzoli and Jahn, 2007)). If the pore expands beyond a certain diameter, the vesicle collapses and integrates into the surface, spilling out its cargo, and the added membrane is retrieved by endocytosis at, or near, the site of exocytosis (i.e. full collapse(Rizzoli and Jahn, 2007)). Hence, fusion pore dynamics define the release kinetics, and the mode of exocytosis and vesicle recycling.

Several studies from flies to mammals have demonstrated that LSV exocytosis differs from exocytosis of smaller vesicles. When LSVs fuse with the apical surface they recruit an actomyosin meshwork that contracts on the LSV membrane and extrudes the cargo (Valentijn et al., 2000; Sokac et al., 2003; Turvey and Thorn, 2004; Nemoto et al., 2004; Yu and Bement, 2007a; Segawa and Yamashina, 1989; Nightingale et al., 2011, 2012; Tran et al., 2015; Rousso et al., 2016; Yu and Bement, 2007b). We have shown that actomyosin contractility on the LSV membrane squeezes out the content without integrating the vesicle into the apical surface (Kamalesh et al., 2021) (Fig. 1 A). Consequently, the vesicle neither collapses into nor detaches from the surface, suggesting that LSVs utilize a distinct mode of exocytosis (which we termed “membrane crumpling”). We have also shown that diffusion between the vesicle and apical membranes becomes restricted after fusion, thereby maintaining apical membrane composition. These observations suggested that membrane homeostasis in exocrine cells is maintained by mechanochemical sequestration of the LSV membrane (Kamalesh et al., 2021) (Fig. 1 A).

**Fig. 1.**
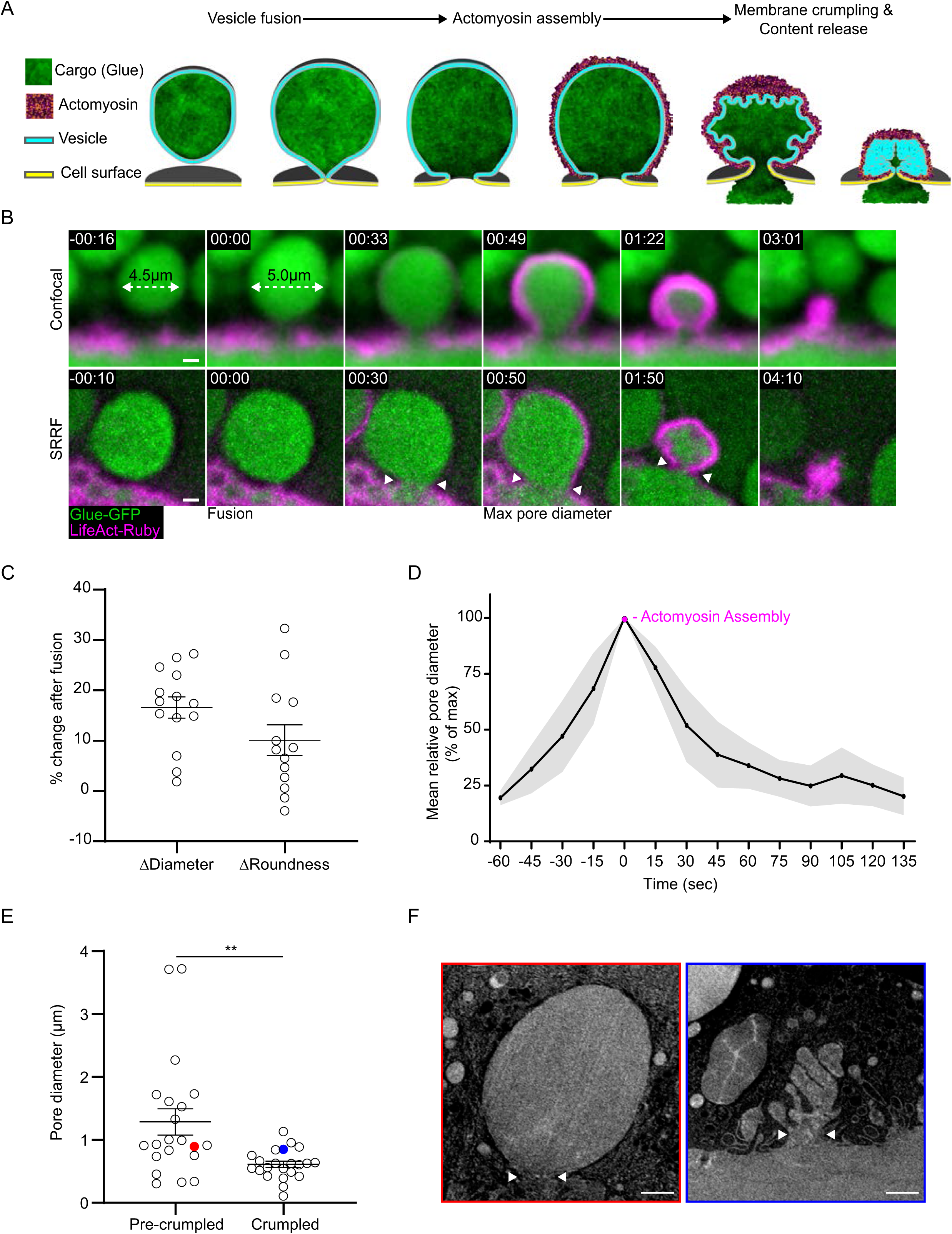
| The fusion pores of LSVs expand and constrict. (A) Schematic representation of LSV exocytosis by vesicle membrane crumpling (MC) – actomyosin is recruited to LSVs after fusion to mediate MC and content release. (B) Time-lapse sequence of representative exocytotic events (Confocal, Top; SRRF, Bottom). Fusion onset is detected by LSV swelling (White dashed arrow; **Fig. S1 A**). Actomyosin recruitment is first visible ∼50 seconds after fusion and followed by content release. The fusion pore expands before actomyosin recruitment and constricts during content release (White arrow heads, Bottom). Time mm:ss, relative to fusion. Glue-GFP (Green) and LifeAct-Ruby (Magenta; expression driven by c135-GAL4). (C) Quantification of LSV swelling as % change in LSV diameter and roundness after fusion (n=14). (D) Mean relative pore diameter over time. Time 0 is at maximal pore diameter (n=12; Gray – SEM). (E) Pore diameter of fused vesicles visualized by FIB-SEM before and after crumpling (n(pores)= 21 and 22 respectively; P: **=0.0026. two-tailed unpaired t test). Red and blue circles denote the individual LSVs shown in (F). (F) Representative slices from FIB-SEM volumes of LSVs before (Red) and after (Blue) crumpling. White arrow heads show the widest plane of the fusion pore where it was measured. Whiskers in (C) and (E) show mean and SEM. Scale bars in (B) and (F) 1µm.

To elucidate the mechanism of exocytosis by membrane crumpling (MC), we used the *Drosophila melanogaster* larval salivary gland (SG) as a model for LSV exocytosis (Biyasheva et al., 2001; Rousso et al., 2016; Kamalesh et al., 2021). The cargo of SG vesicles are adhesive mucinous glycoproteins nicknamed “Glue”, which are used by the fly larva to attach to a solid surface before metamorphosis. Recruitment of active (GTP-bound) Rho1 to the vesicle membrane after fusion triggers two parallel events that are essential for actomyosin meshwork formation (Rousso et al., 2016). Activation of Diaphanous (Dia), a member of the formin family of actin nucleators, which generates linear actin strands that are visible within ∼30 seconds after fusion, and activation of RhoA-kinase (ROCK), leading to the phosphorylation of myosin II light chain and activation of myosin motors on the actin network (Shemesh and Kozlov, 2007; Jaffe and Hall, 2005; Rousso et al., 2016; Segal et al., 2018; Kamalesh et al., 2021).

We now find found that after LSVs fuse they remain connected to the apical surface through dynamic fusion pores that expand to diameters larger than 1 µm and constrict back to 100 nm or less (Kamalesh et al., 2021; Rousso et al., 2016). Utilizing pharmacological and genetic perturbations, we demonstrate that branched actin is essential for pore expansion and stabilization. Conversely, myosin II is required for pore constriction. Under conditions where the fusion pore was destabilized, vesicle content release took place even in the absence of actomyosin, underscoring the presence of the future pore as the cause for the employment of actomyosin contractility during content release. Genetic screening identified several conserved Bin-Amphiphysin-Rvs homology (BAR) domain proteins that control fusion pore dynamics and revealed that the I-BAR containing protein Missing-in-Metastasis (MIM) is essential for pore expansion and stabilization in a dose dependent manner. MIM localizes to the fusion site on the vesicle before fusion and remains associated with the fusion pore throughout secretion. The I-BAR domain is essential but not sufficient for MIM localization and function. Collectively, our results suggest that LSV fusion pore behavior is tightly regulated by BAR domain proteins that act in-concert with actomyosin and branched actin polymerization to facilitate exocytosis, while maintaining apical membrane homeostasis. Moreover, they demonstrate that exocytosis by MC is a distinct mode of exocytosis that depends on the structure and dynamics of the fusion pore.

## Results

### The fusion pores of LSVs expand before actomyosin assembly and constrict during content release

To follow the dynamic changes in LSV fusion pore diameter, we visualized individual vesicles during exocytosis in secreting SGs expressing the content marker Sgs3-GFP (Biyasheva et al., 2001) (Glue-GFP) under the endogenous *sgs3* promoter and the F-actin probe LifeAct-Ruby (expressed via GAL4/UAS using the c135-GAL4 driver) as a marker for actomyosin assembly (Fig. 1 B). The time of vesicle fusion was defined by the swelling and rounding of the LSVs, which occurs at the onset of fusion (Breckenridge and Almers, 1987) (Fig. 1, B-C and S1 A). To visualize and quantify changes in pore diameter during exocytosis, we used super resolution radial fluctuation (SRRF) imaging, which improved the resolution and signal to noise ratio (Gustafsson et al., 2016). We found that following fusion the pore expands during 58 ± 5 seconds, reaching a maximum mean diameter of 1.6 ± 0.2 µm (n=12), and subsequently constricts to diameters below the detection limit, roughly under 100 nm. Actomyosin assembly begins when pores reach their maximal diameter, and pore constriction is accompanied by actomyosin contraction and content release lasting altogether 121 ± 9 seconds (Fig. 1, B and D and S1 B).

To directly visualize the fusion pore at different stages of exocytosis, we used Focused Ion Beam Scanning Electron Microscopy (FIB-SEM) and obtained three-dimensional, high-resolution information from the apical areas of secreting SGs (Fig. 1, E and F). We identified fused LSVs and measured their fusion pore diameter. Pore diameter of early fused LSVs, before membrane crumpling (MC), ranged between 0.2 – 3.7 µm with a mean of 1.3 ± 0.2 µm (n=21). In contrast, pore diameter of crumpled LSVs ranged between 0.1 – 1.1 µm with a mean of 0.6 ± 0.04 µm (n=22), consistent with the live imaging data showing that pores expand until actomyosin assembly initiates and subsequently constrict during actomyosin-mediated MC (Fig. 1, E and F). These observations show that the LSV fusion pore follows a typical sequence of expansion and constriction, suggesting that an active mechanism regulates fusion pore dynamics.

### Branched actin nucleation is essential for fusion pore expansion and stabilization

Pharmacological and genetic inhibition of actin polymerization in chromaffin cells, *Xenopus* eggs, pancreatic acinar cells and *Drosophila* SGs influence content release post fusion (Sokac et al., 2003; Ñeco et al., 2004; Nemoto et al., 2004; Larina et al., 2007; Yu and Bement, 2007b; Doreian et al., 2008; Rousso et al., 2016; Shin et al., 2018). To explore the regulatory role of branched actin polymerization in LSV fusion pore dynamics, we treated SGs with a mild dose of the Arp2/3 complex inhibitor CK666 (Ck666^100µM^) or knocked down (KD) Arp3 expression (Arp3^KD^) by an RNA interference (RNAi) construct expressed under UAS control. When the Arp2/3 complex was perturbed by either method, we observed frequent occurrences of fusion pores that failed to expand and stabilize as compared to pores in control, untreated SGs. These alterations in pore dynamics resulted in LSV behaviors which are reminiscent of the kiss-and-run (KAR) and full collapse (FC) canonical modes of exocytosis (Fig. 1 B, 2, A-C and S2, A and B). KAR events are characterized by LSVs that fuse with the apical membrane but subsequently detach. In these events, LSVs will often undergo multiple cycles of fusion (detected by vesicle swelling) and detachment, both in the absence (KAR w/o actomyosin) or the presence of actomyosin recruitment and subsequent contraction (KAR w actomyosin). KAR LSVs that recruit actomyosin are commonly observed to deform so as to “jump-back” into the cytoplasm, strongly suggesting that their fusion pores were very narrow or resealed (Fig. 2, A and B and 2S, A and B). FC events, on the other hand, are characterized by pores that expand irreversibly without stabilizing, leading to rapid and complete opening of the pore, which flattens and integrates the LSV into the apical membrane, releasing the content before actomyosin recruitment (Fig. 2, A and B and S2, A and B). In contrast to wild-type (WT) SGs, we observed significantly more KAR and FC events in Arp3^KD^ SGs (50± 6% and 27 ± 3 % respectively; Fig. 2 C). In CK666^100µM^ treated SGs we also observed significantly more KAR events and an apparent, but non-significant increase in FC events (37 ± 8 % and 19 ± 9 % respectively; Fig. 2 C).

**Fig. 2.**
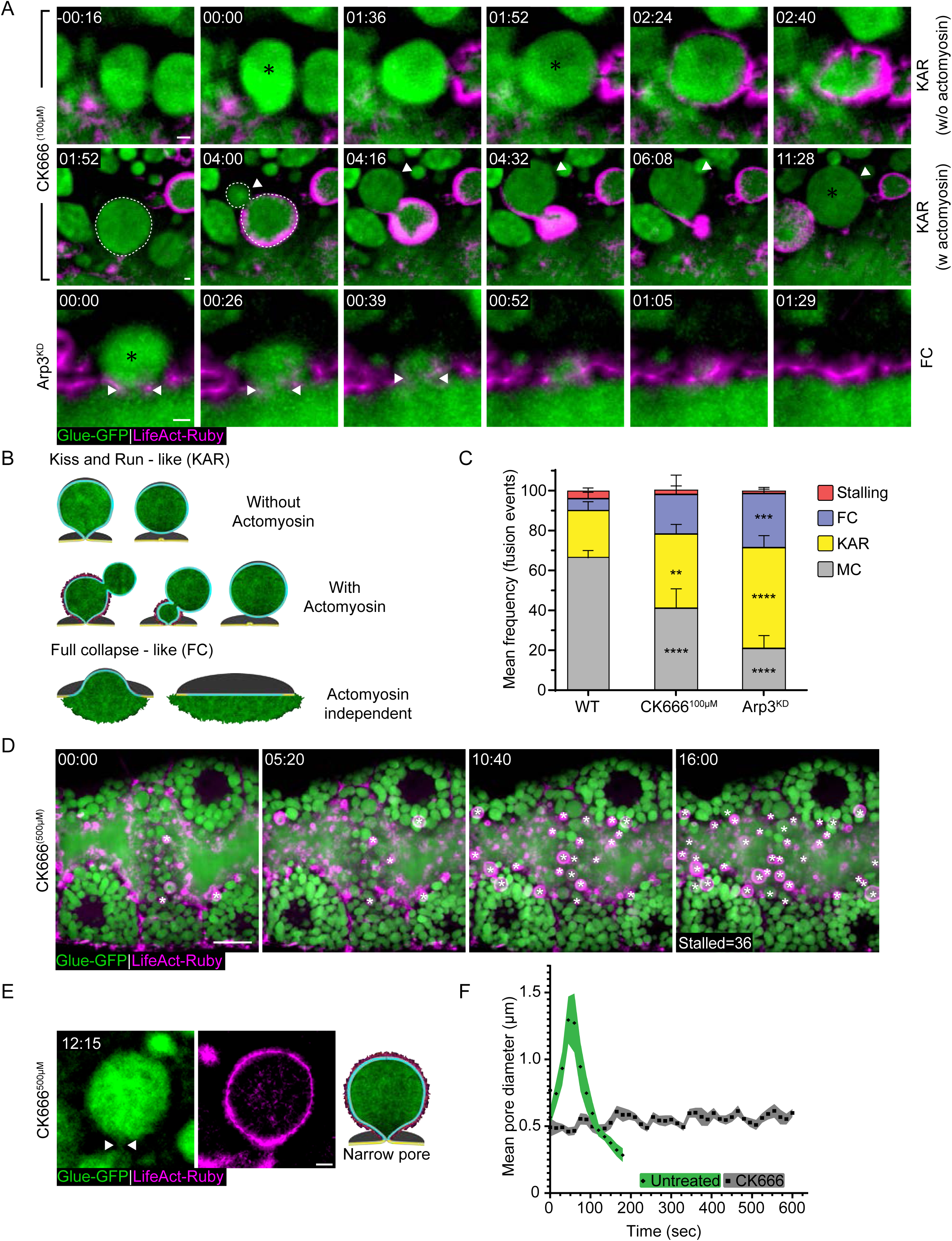
| Branched actin polymerization is essential for pore expansion and stabilization. (A) Time-lapse sequence of representative LSVs from CK666^100µM^ and Arp3^KD^ SGs undergoing KAR and FC (SRRF Z-projection). (Top) KAR w/o actomyosin, appearing as consecutive fusion events (Asterisk) without content release. In this instance, actomyosin was eventually recruited after the second fusion event. (Middle) KAR w. actomyosin, appearing on LSVs that ‘jump back’ away from the apical membrane (Arrowhead and Dashed line). The same LSV fused again at 11:28. (Bottom) FC appearing as content release that precedes actomyosin assembly. The fusion pore expands, and the vesicular membrane is integrated into the apical surface. Glue-GFP (Green) and LifeAct-Ruby (Magenta). (B) Schematic model of KAR (w. and w/o actomyosin), and FC exocytosis (C) Mean frequency (%) of stalling, FC, KAR and MC in WT, CK666^100µM^, Arp3^KD^. Exocytosis by MC is the dominant mode of exocytosis in WT SGs. CK666^100µM^ and Arp3^KD^ present a higher frequency of FC and KAR. Error bars show SEM. Statistical significance with respect to WT frequencies. N(SGs)≥3, n(events)≥200. P values for CK666^100µM^: **(KAR)=0.004044, ****(MC)=0.00002, for Arp3^KD^ ***=0.000199, ****(KAR)=0.000007, ****(MC)<0.000001, two-tailed un-paired multiple t tests corrected using the Holm-Sidak method. (D) Time-lapse sequence of a representative SG treated with CK666^500µM^ (Confocal Z-projection). Stalled, actin coated LSVs (White asterisks) accumulate at the cell apical membrane. Glue-GFP (Green) and LifeAct-Ruby (Magenta). (E) Representative images of stalled LSVs with a narrow pore after CK666^500µM^ treatment. Glue-GFP (Green) and LifeAct-Ruby (Magenta). (F) Change in mean pore diameter over time in control (untreated) and treated SGs showing that while the pore expanded and constricted in control SGs (diamonds; n=12; Green = SEM) it remained stalled with a narrow diameter in treated SGs (CK666 – squares; n=10; Gray= SEM). Fusion = Time 0. Complementary examples to (A) in **Fig. S2 A**. SGs express Glue-GFP and LifeAct-Ruby in (A), (C)-(E). RNAi expression in (A) and (C) under GAL4/UAS control. UAS expression driven in (A) and (C) by *fkh*-GAL4, in (D) and (E) by c135-GAL4. Time mm:ss, in (A), and (E), relative to fusion, in (D) relative to treatment. Scale bars in (A) and (E) 1µm, (D) 20µm.

Treating SGs with a higher dose of CK666 (CK666^500µM^) resulted in the accumulation of stalled, actin coated LSVs at the apical surface of the cell, consistent with previous studies, linking Arp2/3 activity with vesicle contraction (Tran et al., 2015; Rousso et al., 2016) (Fig. 2 D). These LSVs were stalled with narrow pores that did not expand (mean maximal diameter 0.76 ± 0.05 µm, n=10; Fig. 2, E and F), implying that branched actin nucleation through the Arp2/3 complex is, in addition, essential for pore expansion and stabilization. Taken together with the observations of unimpeded content release under MC and FC modes of exocytosis, these results suggest that pore expansion is essential for Glue secretion.

Interestingly, detailed examination of multiple fusion events in WT glands revealed that a significant fraction of these resulted in KAR behavior (23 ± 4%) before the LSVs proceeded to MC (67 ± 3 %; Fig. 2 C and S2 B), and that fusion events occasionally led to FC or stalled LSV behaviors (6 ± 3% and 4 ± 1% respectively). These observations suggest, therefore, that the genetic and pharmacological disruptions exacerbate inefficiencies that are inherent to the LSV secretion process.

### Myosin II is essential for fusion pore constriction

We have previously shown that treating SGs with the Rho-associated protein kinase (ROCK) inhibitor Y-27632, which blocks myosin II recruitment and actomyosin contractility, results in fused but stalled vesicles (Kamalesh et al., 2021; Segal et al., 2018). To explore the role of myosin II in regulating the fusion pore, we examined fusion pore dynamics in these stalled LSVs. We treated SGs with the ROCK inhibitor Y-27632 (100µM,) which induced significant LSV stalling (70% ± 9% of fusion events; Fig. 3 A) and observed that the LSVs were stalled with expanded pores, comparable in diameter to the mean maximal pore diameter of LSVs in untreated SGs (mean max pore diameter 1.8 µm ± 0.09 µm, n=10; Fig. 3, B and C). To elucidate the temporal order of effects on pore dynamics we employed simultaneous treatment with both CK666^500µM^ and Y-27632^100µM^ inhibitors. Under these conditions, SGs presented stalled LSVs with narrow pores (mean maximal diameter of 0.9 ± 0.06 µm, n=9; Fig. 3 C), similar to the effect of the Arp2/3 inhibitor alone. Taken together, these results suggest that myosin II activity is essential for pore constriction downstream of branched actin polymerization, which is essential for pore expansion.

**Fig. 3.**
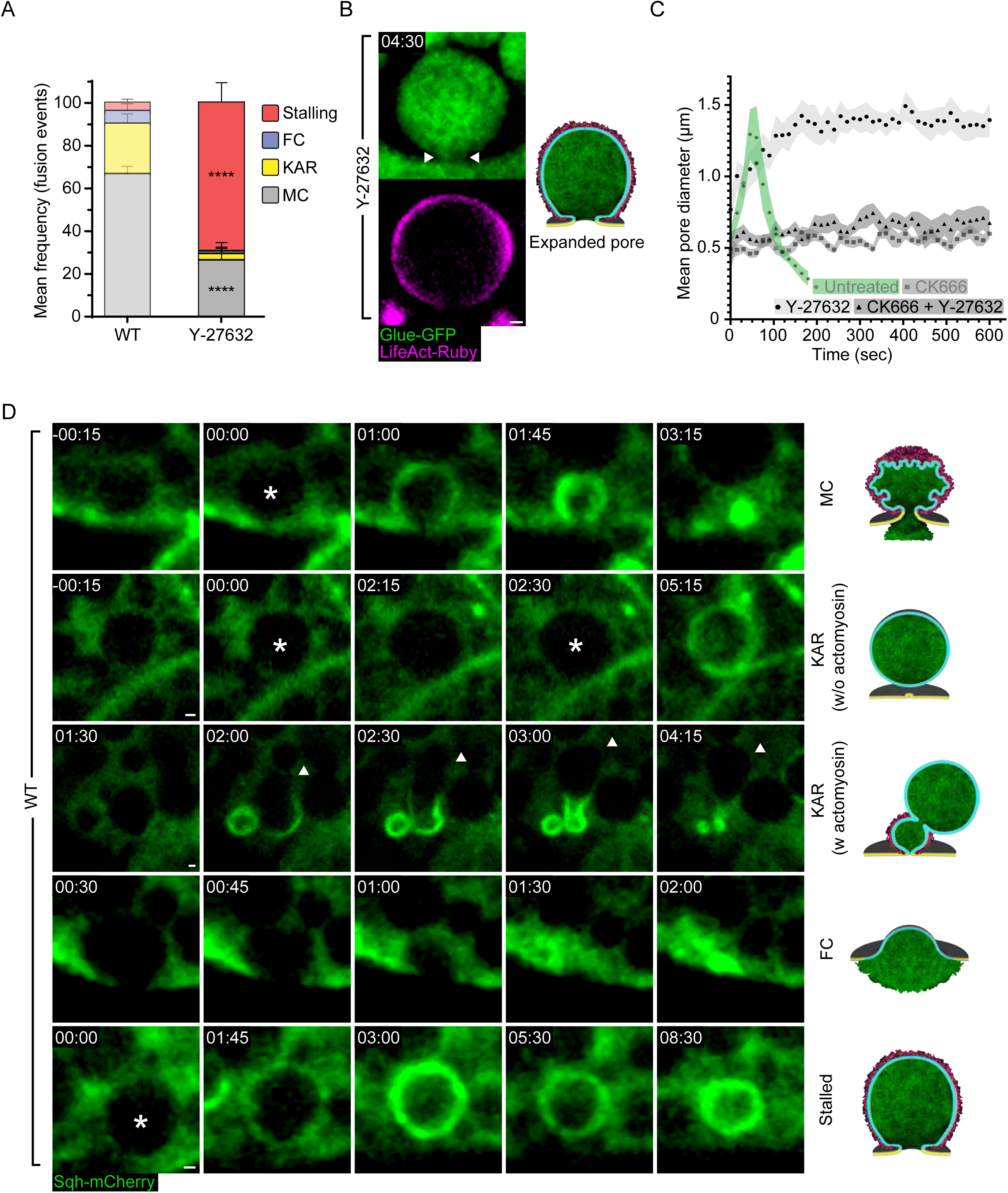
| Myosin recruitment is essential for pore constriction however occurs irrespective of pore dynamics. (A) Mean frequency (%) of stalling, FC, KAR and MC in WT untreated and treated SGs with Y-27632. Exocytosis by MC is the dominant mode of exocytosis in untreated SGs (from **Fig. 2** **C**, semitransparent). Y-27632 treatment resulted in significant LSV stalling. Error bars show SEM. Statistical significance with respect to WT, untreated frequencies. N(SGs)≥3, n(events)≥200. P values: ****(Stalling and MC)<0.000001, two-tailed un-paired multiple t tests corrected using the Holm-Sidak method. (B) Representative images of stalled LSVs with wide pore after Y-27632 treatment. Glue-GFP (Green) and LifeAct-Ruby (Magenta). (C) Change in mean pore diameter over time in control (untreated) and treated SGs showing that while the pore expanded and constricted in control SGs (diamonds; n=12; from **Fig 2** **F** semi-transparent) it remained stalled with a narrow (CK666 – squares; n=10, from **Fig 2** **F** semi-transparent, and CK666+Y-27632 – triangles; n=10) or a wide diameter (Y-27632 – circles; n=9) in treated SGs. Fusion = Time 0. Green and Shades of gray = SEM (D) Time-lapse sequence of representative LSVs undergoing MC (top), KAR (rows 2 and 3), FC (row 4) and Stalling (bottom) in WT SG expressing Sqh-mCherry (Green). LSVs are visible in the background of the cytoplasmic SqhmCherry signal. LSVs that recruit Sqh “jumped back” (KAR w actomyosin; arrowhead). LSVs undergoing FC release their content completely in the absence of Sqh-mCherry signal. Sqh-mCherry was recruited after the LSV membrane integrates into the surface. SGs express Glue-GFP and LifeAct-Ruby in (A-C). UAS expression driven by c135-GAL4. Time mm:ss in (B) and (D) relative to fusion. Scale bars in (B) and (D) 1µm.

To visualize myosin II localization during MC, KAR and FC exocytosis in WT SGs, we used SqhmCherry, a fluorescently tagged variant of the *Drosophila* myosin II light chain homolog *spaghetti squash (sqh)*, expressed under the endogenous *sqh* promoter. (Fig. 3 D). During MC exocytosis, myosin is recruited to the LSVs shortly after F-actin, when the fusion pore is at its maximal diameter (Segal et al., 2018; Kamalesh et al., 2021) (Fig. 1, B and D). MC and content release by actomyosin contractility will then initiate, approximately 60s after fusion (Segal et al., 2018; Kamalesh et al., 2021). The KAR and FC modes of exocytosis were found to be associated with abnormal, albeit informative patterns of myosin II recruitment. During KAR exocytosis, myosin II was only present on LSVs that recruit F-actin and “jump back”, strengthening the notion that the observed LSV deformation is a result of actomyosin contractility applying force on an LSV with a pore that is either narrow or closed. Myosin II was initially absent from LSVs undergoing FC exocytosis but was eventually recruited to the empty flattened LSV membrane that integrated into the apical surface and displayed some contractile activity (Fig. 3 D). These results show that actomyosin recruitment to the LSV membrane can take place regardless of fusion pore behavior, but that pore stabilization is essential for actomyosin mediated MC.

### BAR domain proteins control pore expansion and stabilization

Several proteins have been implicated in the regulation of fusion pore dynamics in small exocytic vesicles including the BAR domain–containing proteins Amphiphysin1, Syndapin2, and Endophilins, which localize to sites of exocytosis (Somasundaram and Taraska, 2018). Since BAR domain proteins associate with membrane curvature, they may represent putative regulators of fusion pore dynamics. Hence, we conducted a candidate-based genetic screen, using fly lines bearing transgenic RNAi constructs directed against BAR-domain containing proteins which are significantly expressed in secreting SGs (Fig. S3 A).

We observed that KD of five out of the eleven BAR-domain containing candidate genes we studied, Missing-in-metastasis (MIM^KD^), Amphiphysin (Amph^KD^), Sorting nexin 1 (SNX1^KD^), Sorting nexin 6 (SNX6^KD^) or Cdc42 interacting protein 4 (CIP4^KD^) – resulted in a qualitative increase in the frequency of FC, KAR or both. Interestingly, we also observed a substantial frequency of compound exocytosis events (in which vesicles fuse to a previously fused LSV), which are rarely observed in WT SGs, in SNX1^KD^, SNX6^KD^, Centaurin beta 1A^KD^, and Syndapin^KD^. SH3PX1^KD^, Rho GTPase activating protein at 92B^KD^, Nostrin^KD^ and CG8176^KD^ did not show a pore-related phenotype (Fig. 4 A and S3 B).

**Fig. 4.**
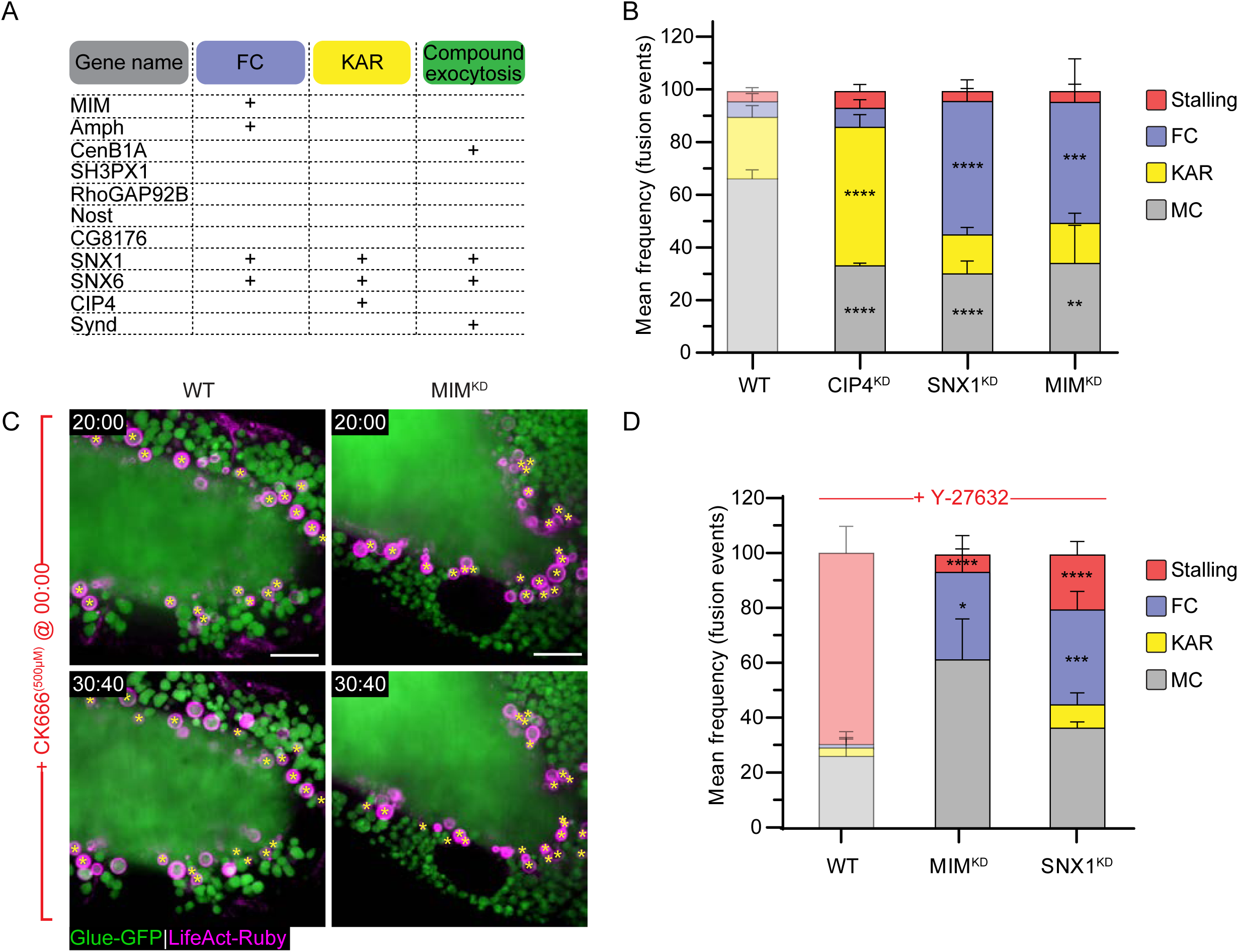
| BAR domain proteins regulate the fusion pore. (A) Qualitative phenotypic RNAi screen for BAR domain containing proteins. MIM^KD^, Amph^KD^, SNX1^KD^ and SNX6^KD^ showed frequent occurrences of FC events, SNX1^KD^, SNX6^KD^, CIP4^KD^ displayed frequent occurrences of KAR events and CenB1A^KD^, SNX1^KD^, SNX6^KD^ and Synd^KD^ showed events of compound exocytosis. (B) Mean frequency (%) of stalling, FC, KAR and MC in CIP4^KD^, SNX1^KD^ and MIM^KD^. CIP4^KD^ resulted in an increase of KAR event frequency, while SNX1^KD^ and MIM^KD^ displayed an increase of FC event frequency. Error bars = SEM. Statistical significance presented is compared to WT SGs. N(SGs)≥3, n(events)≥150. P values for CIP4^KD^: ****(KAR and MC)<0.00001, for SNX1^KD^: ****(FC and MC)<0.000001, for MIM^KD^: ***(FC)=0.000111, **(MC)=0.001267 (two-tailed un-paired multiple t tests corrected using the Holm-Sidak method). (C) Time-lapse sequence of representative CK666 ^(500µM)^ treated WT (left) and MIM^KD^ (right) SGs (confocal Z projection). Glue-GFP (Green) and LifeAct-Ruby (Magenta). Inhibiting the Arp2/3 complex in MIM^KD^ resulted in LSV stalling. Asterisks denotes stalled LSVs. Time mm:ss; relative to CK666 treatment. Scale bars=20µm. (D) Mean frequency (%) of stalling, FC, KAR and crumpling modes in Y-27632 treated WT, MIM^KD^ and SNX1^KD^ SGs. Significantly less stalling was observed, and FC persists at a high frequency in Y-27632 treated MIM^KD^ and SNX1^KD^ SGs. Error bars = SEM. Statistical significance presented is compared to WT treated SGs. N(SGs)≥3, n(events)≥150. P values for MIM^KD^: ****(Stalling)=0.000022, *(FC)=0.011445, for SNX1^KD^: ****(Stalling)=0.000006, ***(FC)=0.000441 (two-tailed un-paired multiple t tests corrected using the Holm-Sidak method). WT and WT Y-27632 treated (from **Fig. 2** **C and 3 A**, semi-transparent) are shown again for comparison convenience in (B) and (D). SGs in (A)-(D) express Glue-GFP and LifeAct-Ruby. RNAi expression under UAS control. UAS-based expression in (A) driven by *fkh*-GAL4, in (B)-(C) by c135-GAL4.

For further analysis, we focused on CIP4, SNX1, and MIM which contain F-BAR, PX-BAR and I-BAR domains, respectively (Fricke et al., 2009; Zhang et al., 2011; Quinones et al., 2010). CIP4^KD^ SGs displayed an increased frequency of KAR exocytotic events (53 ± 4 %), suggesting that CIP4 plays a role in pore expansion (Fig. 4 B and S3 B). In contrast, SNX1^KD^ and MIM^KD^ SGs displayed an increased frequency of FC events, 51 ± 4 % and 46 ± 16 % respectively, suggesting that SNX1 and MIM are important for pore stabilization (Fig. 4 B and S3 B). Consistent with this observation, treating MIM^KD^ SGs with CK666^500µM^ rescued the FC phenotype by stalling the LSVs with a narrow pore similar to the effect of CK666 alone, indicating that pore expansion precedes pore stabilization (Fig. 2 E and 4 C). In contrast, treating MIM^KD^ or SNX1^KD^ SGs with Y-27632 did not alter their FC phenotype, demonstrating that myosin II is required for pore constriction downstream of BAR domain proteins, which are essential for pore stabilization at wide diameters (Fig. 3 B and 4 D). Furthermore, when the pore failed to stabilize, as in MIM^KD^ and SNX1^KD^, vesicle content release was completed even in the absence of actomyosin (Fig. S3 B). These results show that BAR domain proteins are essential for regulating pore dynamics, controlling pore expansion and stabilization.

### MIM localizes to the fusion site and the pore throughout secretion

To determine the localization of CIP4 and SNX1 we examined larval SGs from fly lines expressing CIP4EGFP (Fricke et al., 2009) under an ectopic UAS promoter, and SNX1-GFP under its endogenous promoter. CIP4-EGFP mostly localized to the apical membrane, and SNX1-GFP faintly labeled the apical and lateral membranes of the cells, as well as the LSV membranes. However, we did not observe an enrichment of either CIP4-EGFP or SNX1-GFP at the fusion pore (Fig. S4, A and B).

To determine the localization of MIM we generated fly lines bearing a functional MIM-Emerald (Fig. S5 B) or MIM-mScarlet variants under an ectopic UAS promoter. Upon expression in SGs we observed that both proteins localized to dynamic and mobile cytoplasmic clusters that often merged or divided into smaller clusters (Fig. 5 A and S4 C). Strikingly, we observed small fluorescent MIM clusters that localized to the space between the vesicle and the apical surface before fusion and remained at or in the vicinity of the fusion pore for the duration of exocytosis (Fig. 5 A and S4 C). To determine whether MIM-Emerald localizes to fusion sites we used correlative confocal and FIB-SEM imaging. We targeted cells that showed MIM-Emerald puncta close to the apical surface in secreting SGs. In these cells MIMEmerald was specifically detected at the vesicle membrane in the region that interacts with the apical cell surface, even before fusion and formation of the fusion pore (Fig. 5 B). Cumulatively, our data suggest that different BAR domain proteins regulate distinct steps in fusion pore dynamics, and that MIM localizes specifically to the fusion site on the vesicular membrane, and subsequently to the fusion pore.

**Fig. 5.**
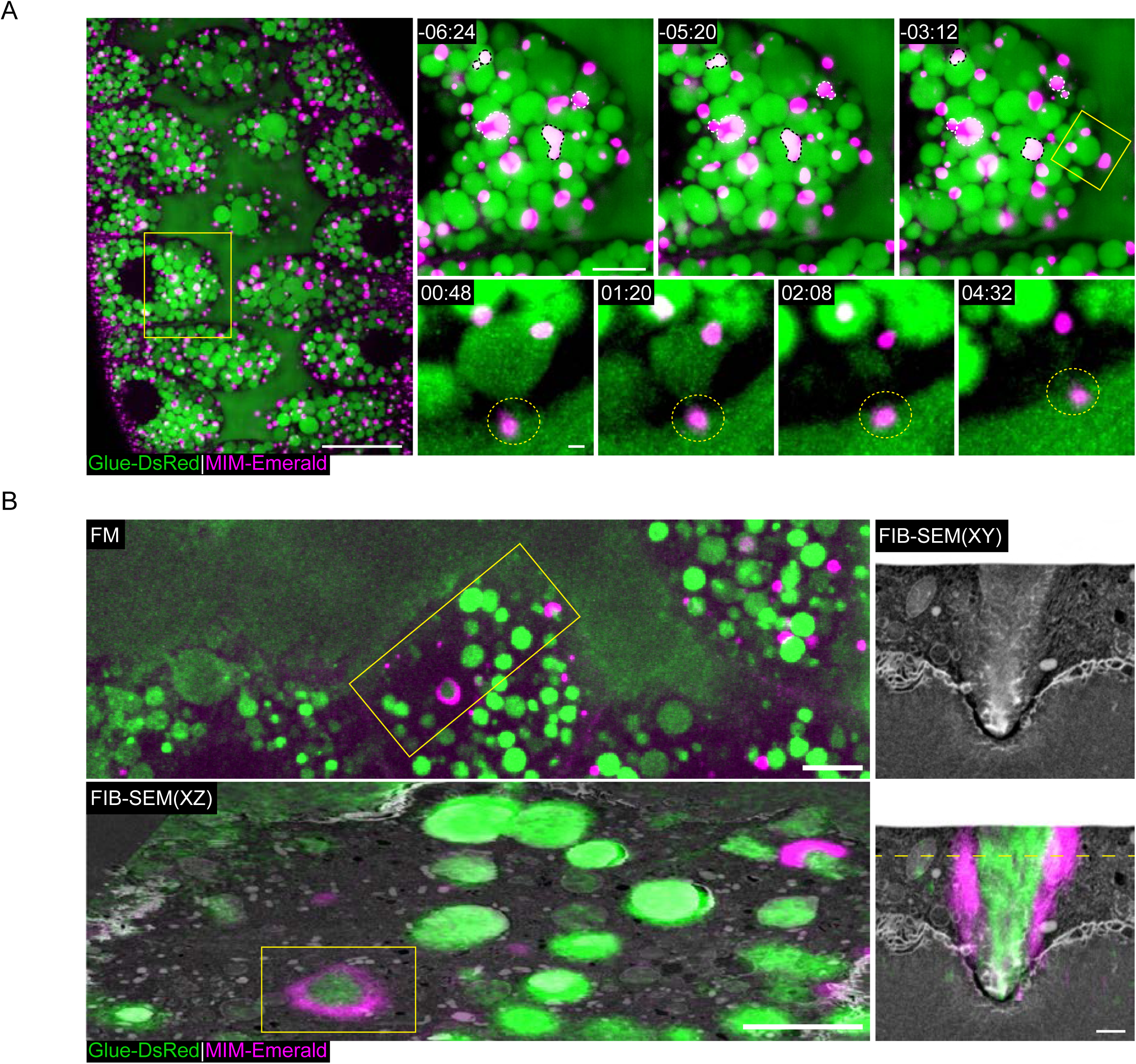
| MIM localizes to the site of fusion and pore formation. (A) Confocal Z projection of a representative SG expressing Glue-DsRed (Costantino et al., 2008) (Green; expressed under the endogenous *sgs3* promoter), and a functional MIM-Emerald (Magenta; **Fig. S5 B**), showing at increased magnification the dynamic localization of MIM to cytoplasmic and apical clusters. Yellow squares mark the magnified area. (Top right) Dynamic MIM cytoplasmic clusters merging (Black dashed lines) and splitting (White dashed lines). (Bottom right) A representative LSV (SRRF Z projection) during exocytosis, showing MIM localization to the sites of fusion and pore formation (Dashed circle) and remaining throughout secretion. Time mm:ss; relative to fusion (of bottom right LSV). Scale bars= 20µm (Left), 10µm (Top right), 1µm (Bottom right). (B) Correlative confocal and FIB-SEM showing MIMEmerald fluorescence associated with the membrane of a putative fusion site. (Top left) Confocal Z projection showing the fluorescence from a representative SG after resin embedding. Yellow squares mark the targeted region for FIB-SEM imaging. (Bottom left) Overlay of the transformed fluorescence microscopy (FM) image onto a resliced XZ plane of the FIB-SEM stack. Correlation precision can be evaluated from the correspondence between the Glue-DsRed (Green) and the LSVs in FIB-SEM. (Right) MIM-Emerald, which appears as a ring in the FIB-SEM XZ plane, as depicted in the XY plane, showing MIM localization to the putative fusion site. Plane shown in the XZ plane (Dashed line). Scale bars= 20µm (Left), 1µm (Middle and Right). UAS expression in (A) and (B) was driven by c135-GAL4.

### MIM is essential for pore expansion and stabilization in a dose dependent manner

To further elucidate the role of MIM in pore regulation, we visualized pore dynamics under progressive MIM loss-of-function conditions. To this end we crossed the chromosomal deficiency line Df(2R)BSC260, bearing a large deletion that includes the MIM gene locus (MIM^Def^) with WT flies, to generate SGs with a single functional copy of MIM (MIM^+/Def^). We did not observe a significant variation in exocytosis in MIM^+/Def^ SGs compared to WT SGs, showing that one copy of the MIM gene is sufficient for exocytosis by MC (Fig. 6 B). Next, we crossed the MIM^Def^ line with a MIM null allele wherein exons 3-10 are deleted (Quinones et al., 2010) (MIM^Null^; Fig. 6 A), to generate a complete MIM null background (MIM^Null/Def^). We observed a significant increase in KAR events (49 ± 3 %; Fig. 6, B and C). The high frequency of KAR exocytosis in MIM^Null/Def^ implies that in the complete absence of MIM, pores fail to expand. These results, taken together with the observation that MIM^KD^ resulted in a higher frequency of FC exocytosis, suggest that MIM is required both for pore expansion and pore stabilization, in a dose dependent manner. In the absence of MIM (MIM^Null/Def^) pores fail to expand, but when if MIM is present at low levels (MIM^KD^) pore expansion can occur, but pores fail to stabilize.

**Fig. 6.**
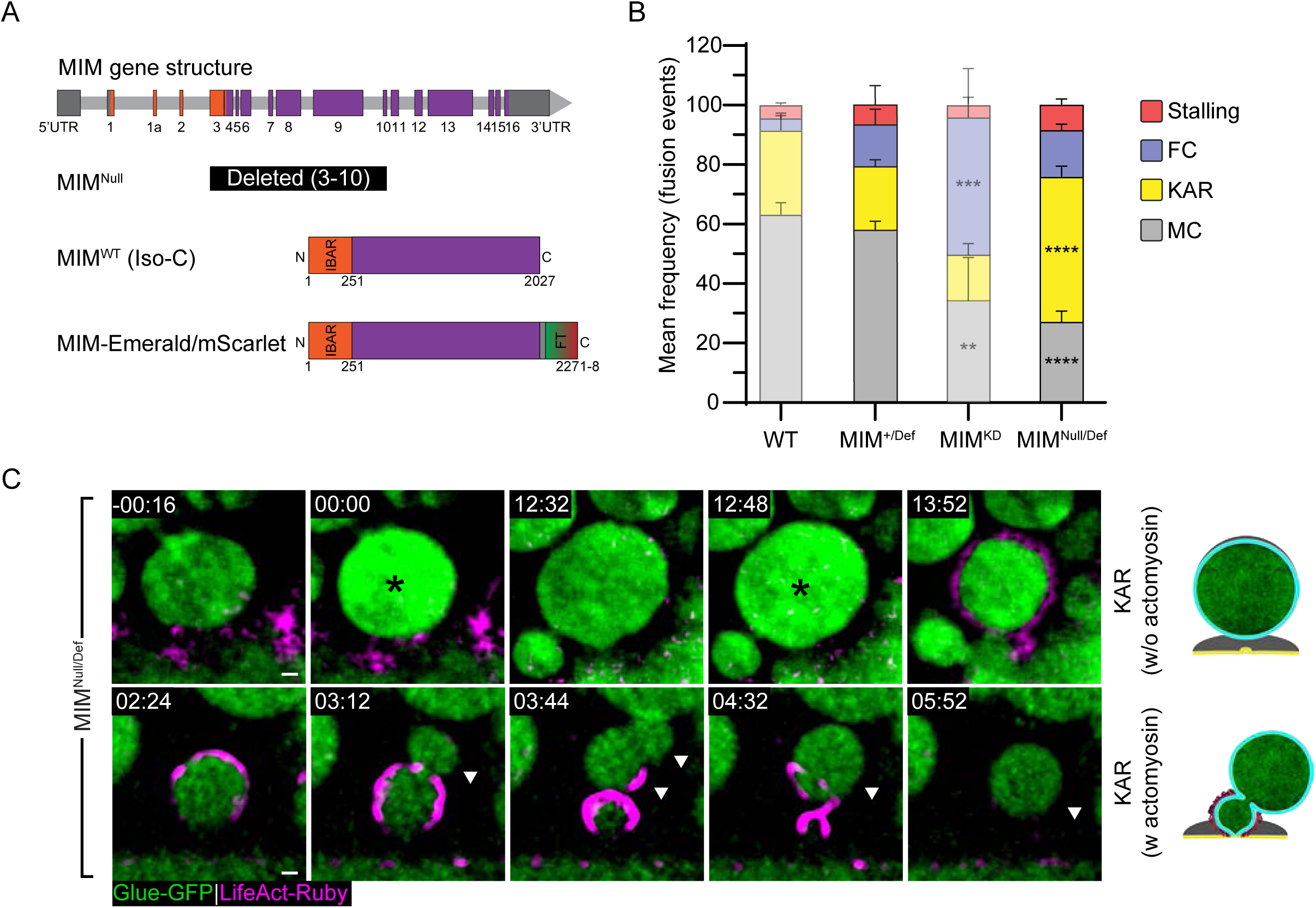
| MIM is essential for pore expansion and stabilization. (A) Schematic of the MIM gene structure and constructs expressed in transgenic lines. Exons (Rectangles), introns (Light gray bars; not to scale), UTRs (Dark ray), Regions encoding for the I-BAR domain (Orange). The null allele MIM^null^ bears an internal deletion of exons 3-10. Isoform C (Iso-C) is the longest isoform and was used to generate MIM-Emerald/mScarlet; Fluorescence tag (FT). (B) Mean frequency (%) of stalling, FC, KAR and MC in MIM^+/Def^, MIM^Null/Def^ and MIM^KD^ (from **Fig. 4** **B**, semi-transparent; presented for convenience) compared to WT (from **Fig. 2** **C**, semi-transparent; presented for convenience). MIM^+/Def^ is not significantly different from the WT. MIM^Null/Def^ resulted in an increase of KAR event frequency as opposed to MIM^KD^, which resulted in an increase of FC event frequency. SG expressing Glue-GFP and LifeAct-Ruby. Error bars = SEM. N(SGs)≥3, n(events)≥200. Statistical significance presented is compared to MIM^+/Def^ SGs. P values for MIM^Null/Def^: ****(KAR)=0.000031, ****(MC)=0.00001 (twotailed un-paired multiple t tests corrected using the Holm-Sidak method). (C) Time-lapse sequence (SRRF Z projection) of representative LSVs from MIM^Null/Def^ undergoing KAR in SGs expressing the Glue-GFP (Green) and LifeAct-Ruby (Magenta) markers. (Top) Consecutive fusion events of the same LSV at 00:00 and 12:48 (KAR w/o actomyosin; Asterisks). (Bottom) LSV undergoing KAR that recruited actin “jumps back” (KAR w actomyosin; arrowhead). Time mm:ss; relative to fusion. Scale bars=1µm. UAS expression in (B) and (C) driven by *fkh*-GAL4.

### The MIM I-BAR domain is essential but not sufficient for MIM localization and function

Previous studies have shown that the I-BAR domain by itself is sufficient to induce membrane remodeling in cells (Saarikangas et al., 2009; Nishimura et al., 2021; Tsai et al., 2022). We therefore aimed to explore the contribution of this domain to MIM function and localization at the fusion pore. To test whether the I-BAR domain is essential for localization and function of MIM, we disrupted it by inserting an EGFP expression cassette in-frame into the I-BAR domain genomic sequence (MIM^ΔIBAR^; Fig. 7 A). We observed that SGs exclusively expressing MIM^ΔIBAR^ (MIM^ΔIBAR/Def^) phenocopied MIM^Null/Def^ and displayed a high frequency of KAR events (68 ± 7%), indicating that the I-BAR is essential for function (Fig. 7, B and C and S5 A). In contrast, SGs carrying MIM^ΔIBAR^ across from the wildtype MIM allele (MIM^ΔIBAR/+^) phenocopied the MIM^KD^ phenotype with a high frequency of FC events (50 ± 13 %; Fig. 7, B and C). Given that a single copy of MIM is sufficient for function (Fig. 6 B), the high frequency of FC events observed for MIM^ΔIBAR/+^ suggests that MIM^ΔIBAR^ has a dominant negative (DN) effect over the wildtype MIM protein. Myosin II is still recruited to the LSV membrane after FC in MIM^ΔIBAR/+^ (Fig. S5 C and 3 D), underscoring the notion that pore stabilization by MIM is not essential for actomyosin recruitment and contractility. These results reinforce the conclusion that MIM activity is required in a dose dependent manner for pore expansion and subsequently for pore stabilization, and show that the I-BAR domain is essential for these functions. Importantly, overexpression of MIM-Emerald in both MIM^Null/Def^ and MIM^ΔIBAR/Def^ SGs rescued the loss-of-function phenotype (Fig. S5 B), strengthening the conclusion that MIM localizes to the fusion pore, as observed with MIM-Emerald, and that the observed changes in pore dynamics are a direct consequence of MIM loss-of-function.

**Fig. 7.**
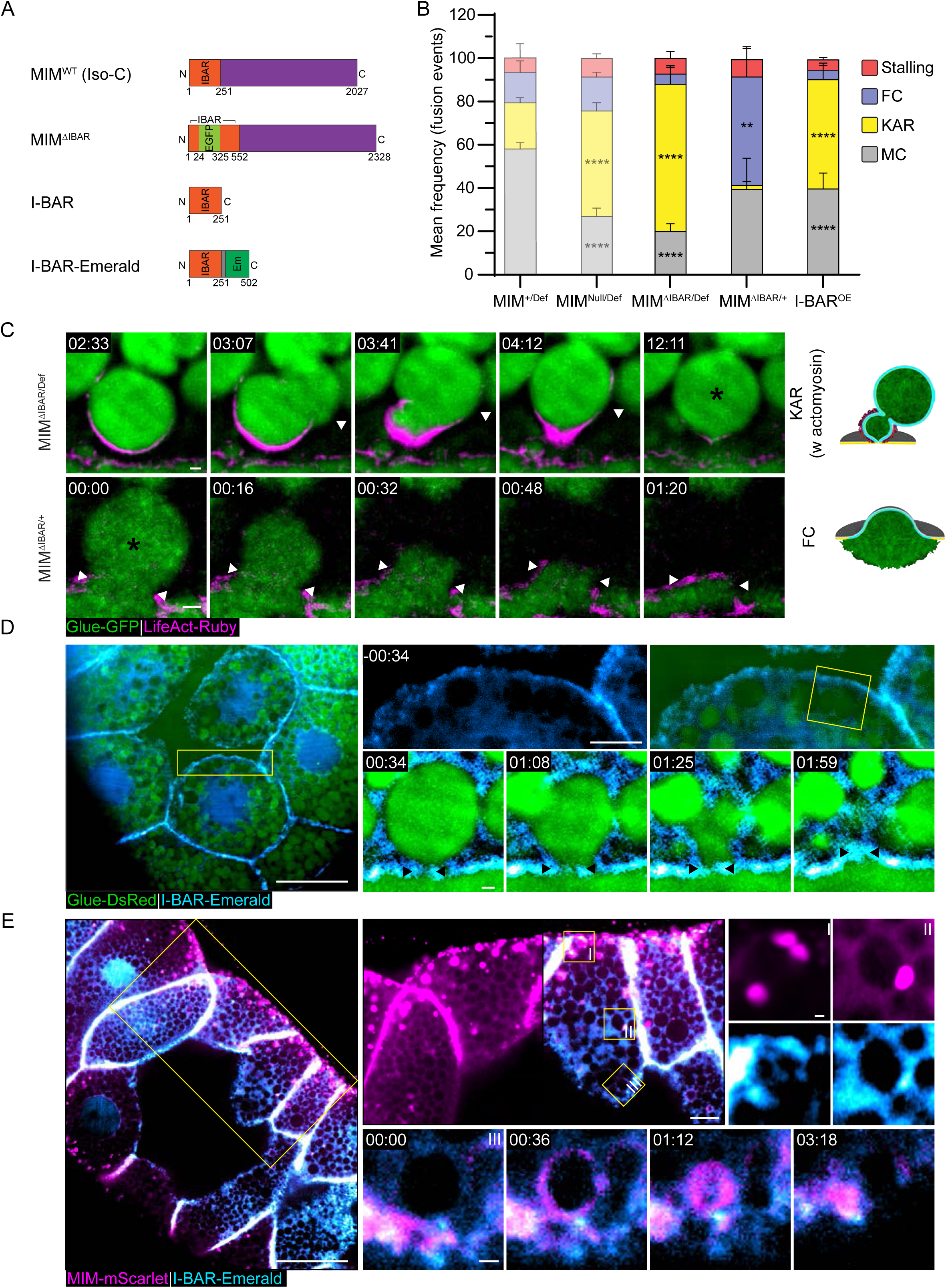
| The MIM I-BAR domain is essential but not sufficient for function and localization. (A) Schematic of the constructs expressed in transgenic lines. MIM^ΔIBAR^ – in-frame knock-in of an EGFP expression cassette into the I-BAR domain. MIM isoform C is illustrated, but all isoforms include the insertion. OE constructs of I-BAR and I-BAR-Emerald are under UAS control. We could not detect MIM^ΔIBAR^ EGFP signal our imaging experiments, most likely as it is under our detection limit (over background). (B) Mean frequency (%) of stalling, FC, KAR and MC in MIM^ΔIBAR/Def^, MIM^ΔIBAR/+^ and I-BAR^OE^ compared to MIM^+/Def^ and MIM^Null/Def^ (from **Fig. 6** **B**, semi-transparent; included for convenience). MIM^ΔIBAR/Def^ phenocopied MIM^Null/Def^ with a high frequency of KAR events. MIM^ΔIBAR/+^ displayed a high frequency of FC events, phenocopying MIM^KD^. I-BAR^OE^ displayed a high frequency of KAR. SGs express Glue-GFP and LifeAct-Ruby. Error bars = SEM. Statistical significance with respect to MIM^+/Def^ for the MIM^ΔIBAR^ expressing SGs and to WT SGs for the I-BAR^OE^ SGs. N(SGs)≥3, n(events)≥150. P values for MIM^ΔIBAR/Def^: ****(KAR)<0.000001, ****(NC)=0.000012, for MIM^ΔIBAR/+^: **(FC)=0.003965, for I-BAR^OE^: ****(KAR)=0.000011, ****(MC)=0.000015 (two-tailed un-paired multiple t tests corrected using the Holm-Sidak method). (C) Time-lapse sequence (SRRF Z projection) of representative LSVs undergoing KAR w. actomyosin and FC (complementary examples in **Fig. S5 A**) from MIM ^ΔIBAR/Def^ and MIM^ΔIBAR/+^ SGs. Glue-GFP (Green), LifeAct-Ruby (Magenta). Fusion events (Asterisks). LSVs that recruited actin “jumps back” (KAR w actomyosin; arrowhead). Expanding pore (FC; Double arrowheads). Scale bars = 1µm. (D) Confocal Z projection of a representative SG expressing Glue-DsRed (Costantino et al., 2008) (Green) and I-BAR-Emerald (Cyan). (Left) Overview of the SG showing the I-BAR-Emerald did not localize to cytoplasmic clusters. (Top right) View of the apical surface showing homogenous I-BAR-Emerald localization on the apical membranes. (Bottom right) Time-lapse sequence of a representative LSV (SRRF Z projection) during exocytosis, showing that I-BAR-Emerald did not localize specifically to the fusion pore (arrowheads). (E) Confocal Z projection of a representative SG ectopically expressing I-BAR-Emerald (Cyan) and MIM-mScarlet (Magenta). (Left) Overview of the SG. (Top right) Magnified views showing that the I-BAR-Emerald and MIM-mScarlet co-localized in cytoplasmic clusters and on the apical membrane. (Bottom right) Time-lapse sequence of a representative LSV during exocytosis showing the IBAR-Emerald and MIM-mScarlet co-localizing at the fusion site and the fusion pore during secretion. UAS expression in MIM^+/Def^, MIM^Null/Def^ and MIM^ΔIBAR/Def^ SGs driven by *fkh*-GAL4, in I-BAR^OE^, (D) and (E) driven by c135-GAL4. In (D) and (E) Yellow squares mark the magnified area. Time mm:ss; relative to fusion in (C-E). Scale bars = 20µm (Left), 10µm (Top right), 1µm (Bottom right).

To determine whether the I-BAR domain is sufficient for MIM function, we generated transgenic flies overexpressing the I-BAR domain in a WT background (I-BAR^OE^; Fig. 7 A). We observed that I-BAR^OE^ SGs have a significantly elevated frequency of KAR events (50 ± 7 %; Fig. 7 B and S5 A), indicating that overexpression of the I-BAR domain alone has a DN effect over the MIM^WT^ protein, and suggesting that the I-BAR domain is not sufficient for MIM function on its own.

To test whether the I-BAR domain is sufficient for MIM localization, we generated flies overexpressing a fluorescently tagged version, I-BAR-Emerald (Fig. 7 A). We observed that unlike MIM-Emerald and MIM-mScarlet, I-BAR-Emerald did not localize to cytoplasmic clusters and did not accumulate in the vicinity of fusion sites. Instead, I-BAR-Emerald was uniformly distributed on the apical and lateral membranes and in the cytoplasm (Fig. 7 D), suggesting that the I-BAR domain by itself is not sufficient for MIM localization. To test for mutual effects on localization that may result from the formation of a complex between the I-BAR domain and full length MIM, we coexpressed I-BAR-Emerald with MIM-mScarlet. The two proteins colocalized in cytoplasmic clusters, in the vicinity of fusion pores, and on secreting LSVs (Fig. 7 E). In addition, MIM-mScarlet localized uniformly on the apical membrane, mimicking the localization of the I-BAR domain (Fig. 7 E). These experiments show that the I-BAR domain can interact with the full length MIM-mScarlet to produce both WT localization and the localization of the I-BAR domain alone. Collectively, these results suggest that the I-BAR domain is essential but not sufficient for MIM localization and function.

## Discussion

Secretory vesicle exocytosis is a fundamental biological process of great physiological importance. Secretory vesicle size differs between neuronal, endocrine, and exocrine cells, and correlates well with their content and physiological function. Exocrine cells tend to generate large secretory vesicles (LSVs) that are at least an order of magnitude larger than conventional vesicles, sometimes reaching the size of a yeast cell. As vesicles increase in size, their surface area-to-volume ratio decreases. Hence, LSVs represent the most economical way to package large quantities of cargo. The larger size of LSVs may also facilitate packaging of viscous macromolecular mixtures such as tears, saliva, surfactants, and adhesives such as Glue. However, increasing the size of secretory vesicles poses significant challenges to vesicle biogenesis, trafficking, fusion, content release, recycling, and maintenance of the limited apical cell surface.

We analyzed fusion pore dynamics of LSVs, using the *Drosophila* larval salivary glands as a model for exocrine tissues. We observed that when LSVs fuse to the apical surface, the fusion pore expands rapidly, stabilizes with a wide diameter, and subsequently constricts to diameters that fall below the diffraction limit of light (Fig. 1 and S1). The ability to arrest the pore at distinct stages suggested that an intricate molecular machinery controls pore expansion, stabilization and constriction during LSV exocytosis. Eliminating branched actin polymerization, MIM, and CIP4 inhibits pore expansion, resulting in pore resealing and kiss-and-run like (KAR) exocytosis (Fig. 2, 4, 6-7, S2, S3 and S5). On the other hand, KD of branched actin polymerization, MIM, SNX1, SNX6 and Amph inhibits pore stabilization, resulting in irreversible pore expansion and full collapse-like (FC) exocytosis (Fig. 2, 4, S2 and S4). Inhibiting myosin II recruitment, arrests pore constriction and membrane crumpling (MC) without effecting pore stabilization and results in vesicle stalling with wide pores that fail to release their cargo (Fig. 3).

Consistently, expanded fusion pores, often referred to as omega (Ω)-shaped profiles, also persist for several minutes in pancreatic acinar cells and alveolar type II cells (Nemoto et al., 2001; Haller et al., 2001). Similarly, fusion pores during secretion of Von Willebrand factor from Weibel-Palade bodies also limit vesicular membrane integration, allowing endocytosis to occur specifically from the vesicular membrane even after the pore reseals (Stevenson et al., 2017). Taken together, these observations suggest that maintaining membrane homeostasis via pore regulation is a conserved mechanism during LSV exocytosis.

We propose that MC exocytosis is a distinct mode of exocytosis, which requires dedicated protein modules to expand, stabilize and constrict the fusion pore. Fusion pore stabilization enables actomyosin recruitment to the LSV membrane such that during actomyosin contraction, the vesicle membrane is folded without incorporating and diluting the apical surface (Fig. 8). Thus, apical membrane homeostasis is maintained during secretion. When the fusion pore fails to stabilize actomyosin is still recruited, but only after the content is released by FC, and at the expense of maintaining membrane homeostasis (Fig. 8). These results show that exocytosis *per se* and myosin II recruitment can be uncoupled, and that actomyosin is only essential for LSV exocytosis if the fusion pore is stabilized by a dedicated protein machinery (Kamalesh et al., 2021). Moreover, they show that fusion pore stabilization and not actomyosin, physically sequesters the vesicular membrane during exocytosis.

**Fig. 8.**
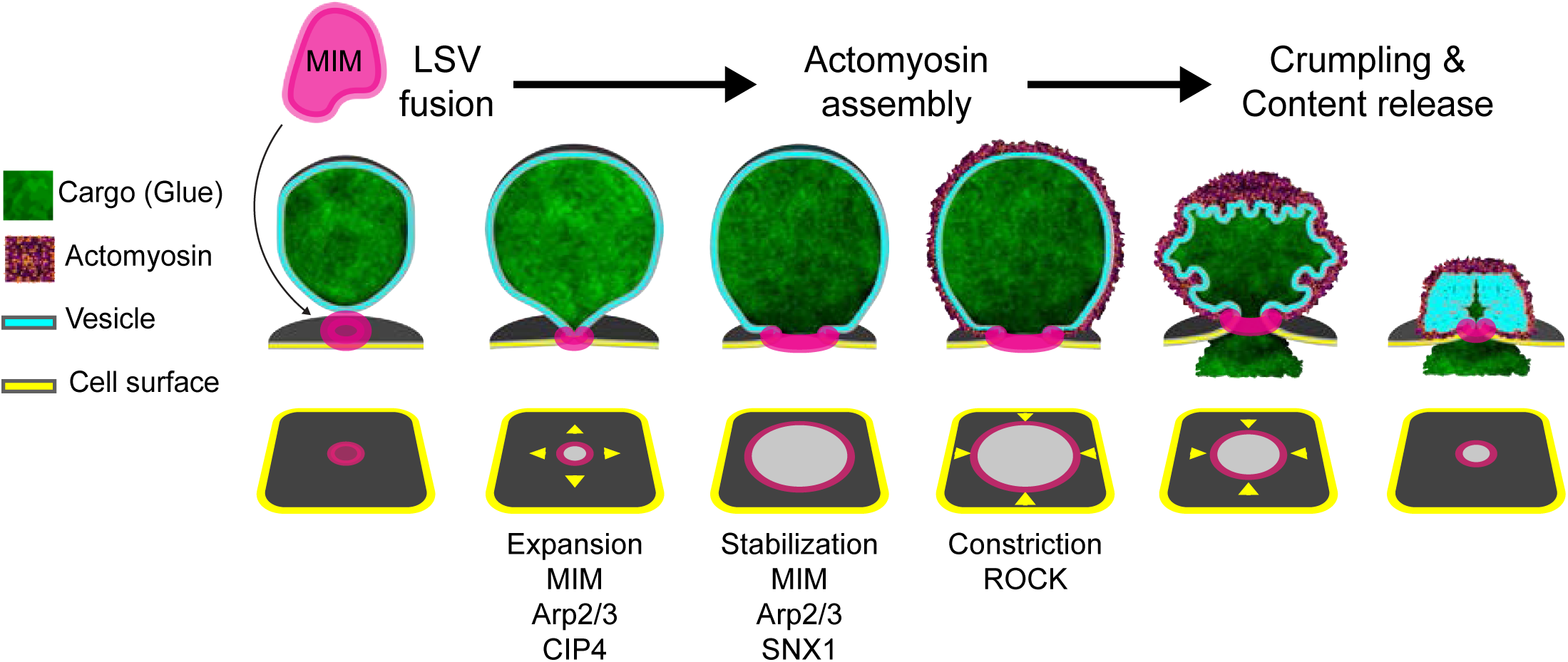
| BAR domain proteins, actin polymerization and myosin II control fusion pore dynamics to facilitate crumpling exocytosis. Schematic model of pore expansion, stabilization and constriction as distinct steps in a sequence that facilitates exocytosis by actomyosin-mediated MC. The pore is regulated at each step by distinct components which include branched actin polymerization, myosin II and BAR domain proteins. The I-BAR protein MIM cooperates with actin and myosin to control fusion pore dynamics of large secretory vesicles. MIM localizes to the future fusion site on the vesicle. After fusion, the pore expands in an Arp2/3, MIM and CIP4 dependent manner. The pore stabilizes with a wide diameter in a MIM dependent manner, preventing full collapse and membrane integration. SNX1 is also essential for efficient pore stabilization. Pore constriction depends on myosin II and initiates during actomyosin mediated MC. Orchestrated dynamics of the fusion pore is essential for MC and insulation of the apical cell membrane during exocytosis.

Actomyosin recruitment and contractility still occur on the residual LSV membrane, even after its integration into the surface by FC (Fig. 3 and S5). This restricted actomyosin recruitment to the fused vesicle in the absence of a fusion pore indicates that the fusion pore in-of-itself does not chemically insulate the LSV membrane from the apical surface. The lack of mixing between the membranes may thus represent a distinct intrinsic set of properties between the vesicular and apical membranes. In this context, it would be interesting to test whether the vesicular and apical membranes do mix in SNX1^KD^, SNX6^KD^, Centaurin beta 1A^KD^, and Syndapin^KD^ that result in compound exocytosis, which might be attributed to membrane mixing and the acquisition of an apical cell membrane identity by LSVs after fusion.

The formation of dynamic fusion pores that expand, stabilize, and constrict, requires transitions between membrane shapes and curvatures. Such transitions rely on molecules that can sense these changes and respond. Ca^2+^, F-actin and myosin II have been implicated in fusion pore expansion during exocytosis of LSVs (Haller et al., 2001; Larina et al., 2007; Doreian et al., 2008; Ñeco et al., 2008; Bhat and Thorn, 2009) (reviewed in (Miklavc and Frick, 2020)). Consistently, we found that branched actin polymerization and CIP4 are essential for LSV fusion pore expansion (Fig. 2, 4 and S3). Given the CIP4^KD^ phenotype and CIP4 localization (Fig. S4), we speculate that CIP4 might interact with the early fusion pore through its F-BAR domain (Shimada et al., 2007), and influence actin polymerization on the apical membrane through interactions with WASP, WAVE, Arp2/3 activators, or Dia, leading to pore expansion (Fricke et al., 2009; Yan et al., 2013).

We demonstrate a role for the actin cytoskeleton in fusion pore stabilization, as previously hypothesized (Haller et al., 2001). This function is distinct from the activity of actomyosin during vesicle constriction. Actomyosin recruitment and contractility are still observed under mild Arp2/3 inhibition (CK666^100µM^ & Arp3^KD^), supporting the notion that actin fulfils non-overlapping roles at the fusion pore and on the vesicular membrane (Tran et al., 2015; Kamalesh et al., 2021; Bhat and Thorn, 2009) (Fig. 2 and S2). Moreover, pore expansion and stabilization occur prior to actomyosin contraction which is only essential for extruding the cargo through a stabilized and constricting pore. Actomyosin contractility on fused vesicles has been reported in many systems utilizing LSVs, where it might provide the force necessary to extrude the cargo through dynamic fusion pores (Jerdeva et al., 2005; Yu and Bement, 2007b; Miklavc et al., 2011, 2012; Masedunskas et al., 2011; Nightingale et al., 2011; Tran et al., 2015; Miklavc et al., 2015; Kamalesh et al., 2021). Myosin II has also been implicated in pore expansion in pancreatic acinar cells and chromaffin cells utilizing large and medium sized vesicles respectively (Doreian et al., 2008; Ñeco et al., 2008; Bhat and Thorn, 2009). We found however, that myosin II is essential for pore constriction in the *Drosophila* SG (Fig. 3). These differences may arise from variation in the myosin motors employed, their activity or their regulation (Bhat and Thorn, 2009; Rousso et al., 2016; Milberg et al., 2017; Miklavc et al., 2015; Sokac et al., 2006; Yu and Bement, 2007b; Kittelberger et al., 2016; Jerdeva et al., 2005). Nevertheless, the emerging view is that a combination of actin polymerization and myosin motors supply, at least in part, the forces needed to expand, stabilize, and constrict the LSV fusion pore and independently to fold and retain the vesicular membrane while extruding its content.

How are these and other forces directed to act at a specific time, place, and direction in order to control the fusion pore diameter? BAR-domain proteins have direct membrane interacting, shaping, and curvature sensing capabilities, in addition to indirect membrane remodeling activities through interactions with the cytoskeleton (reviewed in (Kozlov et al., 2014; Simunovic et al., 2019)). Indeed, we found that BAR-domain proteins are required for pore expansion and stabilization (Fig. 4 and S3). Specifically, we show that MIM is a key regulator of LSV fusion pore dynamics. MIM localizes to the fusion site prior to fusion, most likely associated with the vesicular membrane. Following fusion, it remains associated with the fusion pore throughout secretion (Fig. 5 and S4). We further show that MIM is important for pore expansion and stabilization in a dose-dependent manner (Fig. 6, 7 and S5) and that the I-BAR domain of the protein is essential but not sufficient for its localization and function (Fig. 7 and S5). Hence, we conclude that MIM fulfills opposing functions by initially promoting pore expansion and subsequently limiting pore dilation.

MIM has both membrane curvature sensing and shaping activities *in-vitro* and *in-vivo* (Saarikangas et al., 2015; Chaudhary et al., 2014; Galbraith et al., 2018; Quinones et al., 2010). The MIM I-BAR domain not only favors negative membrane curvature but can also induce membrane protrusions by directly bending the membrane (Mattila et al., 2007). MIM is also able to interact with positive membrane curvature utilizing amphipathic α helices within the I-BAR domain which are inserted into the membrane (Drin et al., 2007; Bhatia et al., 2009). In addition, MIM harbors a proline-rich domain and a C-terminal WH2 domain, that facilitates interaction with Cortactin (through its SH3 domain) and with the actin cytoskeleton, providing an extensive framework for dynamic membrane remodeling (Quinones et al., 2010; Lin et al., 2005; Mattila et al., 2003; Parker et al., 2023).

The formation of an initial nanometric pore represents an extreme change in membrane curvature. The I-BAR domain acting as a membrane curvature sensor, may activate MIM to recruit branched actin polymerization through the WH2 domain (Quinones et al., 2010; Lee et al., 2007), resulting in pore expansion and recruitment of additional MIM dimers. As the pore expands, membrane curvature decreases and MIM concentration increases, promoting negative curvature on the pore that opposes the forces of actin polymerization, stabilizing the pore prior to pore constriction. Thus, our study suggests a dual function for MIM as a membrane curvature sensor and activator of local actin polymerization at low concentration, and as a membrane shaping protein at high concentration, as previously proposed (Zhao et al., 2011), demonstrating the physiological relevance of such a mechanism. Consistently, MIM plays a similar role in the closure of toxin induced transendothelial cell tunnels by localizing to the rim of the hourglass shaped structure and promoting Arp2/3 mediated actin polymerization (Maddugoda et al., 2011; Fedorov and Shemesh, 2017).

Strikingly, MIM also localizes in *Drosophila* SGs to cytoplasmic clusters/granules that displayed liquid-like behavior, which is a characteristic of protein liquid-liquid phase separation(Brangwynne et al., 2009) (Fig. 5 and S4). *In silico* analysis suggests MIM contains an extended low complexity domain with several putative phosphorylation clusters that are conserved between human and *Drosophila* MIM in addition to the I-BAR domain, indicating a high degree of regulation. In addition, clustering of IRSp53 and MIM was observed *in vitro* and *in vivo* before filopodial elongation (Disanza et al., 2013; Prévost et al., 2015; Saarikangas et al., 2015; Tsai et al., 2022). The role of the low-complexity region in generating the MIM cytoplasmic granules is underscored by the observation that the I-BAR alone mis-localizes to the basolateral membranes, but is targeted to MIM granules in the presence of the full-length protein (Fig. 5 and 7). An interplay between phase separation and assembly on membranes in a curvature-dependent manner, might be part of the mechanism allowing MIM and its interacting partners to pre-assemble the machinery that controls fusion pore dynamics.

Increasing pore size from the nanometric to the micrometric scale necessitates molecules that can assemble and control the expansion, stabilization and constriction of larger fusion pores. Studies in smaller vesicles identified several factors affecting fusion pore dynamics, including Ca^+2^ (Alés et al., 1999; Elhamdani et al., 2006), the SNARE protein complex (Archer et al., 2002; Wang et al., 2003; Wu et al., 2017; Fang et al., 2008; Vardjan et al., 2013; Neuland et al., 2014; Hastoy et al., 2017), dynamin (Graham et al., 2002; Tsuboi et al., 2004), and BAR domain proteins (Llobet et al., 2008; Somasundaram and Taraska, 2018). While some components might be shared across scales, others appear to be unique to LSVs. Our findings sketch the outlines of a dedicated regulatory machinery unique to LSV fusion pores, especially under circumstances where cargo viscosity or membrane homeostasis needs to be addressed. The unique fusion pore of LSVs defines a distinct mode of exocytosis, that consists of fusion pore stabilization followed by actomyosin recruitment and contractility that extrudes the contents. This type of exocytosis assures that the vesicular and apical membranes remain distinct, such that apical membrane homeostasis is maintained, and the vesicular membrane can be retrieved by the slower process of endocytosis (Kamalesh et al., 2021). Modulating fusion pore dynamics results in aberrant KAR and FC types of secretion, highlighting the critical role of the underlying machinery in maintaining the integrity of the secretion process and the health of the secretory tissue.

## Methods

### *Drosophila* strains and rearing conditions

*Drosophila* fly lines obtained from the Bloomington Drosophila Stock Center (NIH P40OD018537), the Vienna Drosophila Resource Center (VDRC, www.vdrc.at) or generated (see below) by this study are summarized in Supplementary Table 1. UAS-CIP4-EGFP (Fig. S4 B; Fricke et al., 2009) was received as a kind gift from Sven Bogdan (Marburg), MIM^null^ (Fig. 6-7 and S5; Quinones et al., 2010) was received as a kind gift from Helen Zenner (Cambridge), Sgs3-DsRed (Fig. 5 and 7; Costantino et al., 2008) was received as a kind gift from Julie Brill (Toronto).

All fly stocks were reared using standard cornmeal, molasses and yeast media at 21°C in a temperature-controlled room. Crosses and flies used from imaging experiments were grown in 25°C incubators without internal illumination. Parent flies producing progeny used for experiments were kept at low density (20-25 flies per bottle) in food bottles and transferred to a new bottle with fresh food every 3-4 days. Imaging experiments were performed on *ex-vivo* cultures of 3^rd^ instar *Drosophila* larval SGs. Larvae from crosses were used without distinguishing between sexes, as no obvious sex specific differences in secretion of SG were observed.

### Generation of transgenic flies

To generate UAS-MIM-Emerald – MIM-Emerald cDNA was synthetized (Genscript). The sequence of MIM Isoform C (longest transcript) was used, with the Emerald sequence flanked by an upstream 15 amino acids (aa) linker and 3 stop codons at the 3’ end. The synthesized cDNA was cloned into pUASt-attB and injected into attP40 (for 2^nd^ chromosome insertion) and attP2 (for 3^rd^ chromosome insertion) flies.

UAS-MIM-mScarlet was generated by replacing Emerald in UAS-MIM-Emerald with mScarlet to generate pUASt-attB-MIM-mScarlet which was injected into attP40 flies (2^nd^ chromosome insertion).

The I-BAR domain was generated by PCR from the MIM isoform C cDNA and cloned into pUASt-attB, injected to attP2 flies (3^rd^ chromosome insertion). MIM was replaced with I-BAR in pUASt-MIM-Emerald to generate pUASt-attB-I-BAR-Emerald which was injected into attP40 flies (2^nd^ chromosome insertion).

To generate MIM^ΔIBAR^, we used the MiMIC system (Venken et al., 2011). Plasmid 1298 (DGRC Stock 1298: pBS-KS-attB1-2-PT-SA-SD-0-EGFP-FIAsH-StrepII-TEV-3xFlag) was injected into MI06553 (BDSC 41450) a MiMIC inserted site between exons 1 and 1a of MIM (location 2R:6947294 [+]), resulting in the EGFP expression cassette translated in all MIM isoforms, inside the I-BAR domain (Fig. 6 A and 7 A). We were unable to detect the MIM^ΔIBAR^ EGFP signal over background, most probably as it was below our detection limit. All embryos microinjections was performed by BestGene Inc.

### Culturing third instar SGs for live imaging

SG culturing was performed as previously described (Rousso et al., 2016). In brief, SGs from third instar larvae were dissected out in Schneider’s medium and transferred to a 35-mm dish, with a 10 mm #1.5 glass bottom well (Cellvis D35-14-1.4-N), containing 100 ml of fresh medium for live imaging. SGs that were naturally secreting were identified by their expanded lumen, visible under a stereomicroscope before imaging. The SGs are visible in the live larvae, such that larvae that were still not in the secreting phase could be returned to the growing bottle and used at a later time. Ecdysone treatment to induce secretion was not used in this study.

### Confocal image acquisition

Imaging of *Drosophila* SGs performed using a Yokogawa automatic Spinning Disk confocal scanning unit (CSU-W1-T2) mounted on an inverted Olympus IX83 microscope. 60x 1.4 NA and 100x 1.49 NA oil immersion objectives were used for data acquisition. Images were captured by dual back illuminated Prime 95B sCMOS cameras (Photometrics), controlled by VisiView software (Visitron Systems GmbH). The following fluorescence excitation and emission filter sets were used: 525/50 nm for GFP, EGFP and Emerald and 609/54 nm for Ruby, mCherry, DsRed and mScarlet. When dual cameras were used, a 561 longpass D2 dichroic mirror was included in the light path before the cameras. Image acquisition was performed using a custom imaging script in Ironpython3 (available upon request). In brief, to produce a single image or a frame in a movie, multiple images were rapidly acquired, which are than either i) projected (intensity projection) to a single image using Fiji (Schindelin et al., 2012), allowing post processing to either improve signal to noise ratio using average projection or to enhance weak signal using sum projection, otherwise not possible on a spinning disk confocal; or ii) processed to super-resolution via the Fiji NanoJ-SRRF plugin (Gustafsson et al., 2016) (see below). Live imaging data used for SRRF contained at least 10 exposure images per single image. To capture the pore (in XZ), no more than 0.5µm was used for slice interval. To capture secretion and pore dynamics, time interval (of final data) was no more than 18 seconds per frame of the time-lapse.

### Super-resolution image processing

Imaging data captured using one of the custom imaging scripts, is made up of multiple image groups, each intended to make up a single image in the processed data. The data is processed using a semi-automated custom Fiji script (available upon request). The script accepts appropriate imaging data folder (or multiple data folders for batch processing), and asks the user to input imaging parameters such as the magnification used, the number of slices and time points or the number of channels and which were used.

The script than prepares a RAW map of the data – projecting the imaging data using the intensity projection Fiji function of Average, Maximum or Sum intensity projection. Users can either use this projected data as is without super resolution or continue and select the time, slice range and area that they wish to be processed into SRRF. The data is then processed into super resolution using the NanoJ-SRRF Fiji plugin calling the plugin individually for each of the final images that will be created, the arguments used in the SRRF plugin are also controlled by user definitions collected by the script. The end result is a ready SRRF, 3D movie, single plane movie or a Z stack in the area, time and slices as chosen by the user. Additional image processing was performed using Fiji for cropping and adjustment of brightness/contrast for visualization purposes only.

### Vesicle 3D segmentation

To demonstrate vesicle expansion upon fusion (Fig. S1 A) we used the surface function in Imaris software. The outlines of the glue signal were marked manually in each plane to create the final surface.

### Drug treatment of SGs

To inhibit ROCK and vesicular secretion, SGs were treated with Y-27632 (100µM final concentration; Sigma-Aldrich) for 20 mins with mild shaking at RT, before imaging (Segal et al., 2018). To inhibit Arp2/3, SGs were treated with CK666 (Sigma-Aldrich); 100µM (mild inhibition) or 500µM (stalling) final concentration. Several treatment protocols have been used depending on the experiment. For quantification of mode of exocytosis distribution (Fig. 2, A and C, 3 A and S2 A), SGs were treated with CK666 (100µM) for 30 minutes with mild shaking at RT, before imaging. To induce vesicle stalling and to measure pores (Fig. 2, E and F and 3 C), SGs were treated with CK666 (500µM) immediately prior to imaging. For double treatment with both Y-27632 and CK666 (Fig. 3 C) SGs were first treated with Y-27632(100µM) for 15 minutes with mild shaking at RT, then CK666 (500µM) was added to the imaging medium and imaging started immediately after. For the MIM KD phenotype under CK666 treatment (Fig. 4 C) SGs were treated with CK666 (500µM) for 20 minutes with mild shaking at RT, before imaging. For all conditions imaging was performed in the presence of the inhibitor/s.

### Quantification of pore diameter

Pore measurements were carried out exclusively on SRRF data, and in the XY plane only. For pores positioned along the XZ plane, the 3 planes surrounding the widest plane of the pore were chosen and projected using maximal or average intensity projection using Fiji (Schindelin et al., 2012). For pores positioned along the XY plane, more planes could be used for the intensity projection as long as the pore in the membrane is clearly visible. Measurements were perform by hand in Fiji by drawing the shortest line across the fusion pore in each frame of the movie. Only pores with clear outlines were chosen for quantifications.

### Quantification of modes of exocytosis frequency distribution

Quantification was done on averaged, bleach corrected (Histogram matching), non-SRRF whole gland time-lapse data sets, using the Cell counter plugin in Fiji (Schindelin et al., 2012). First, fusion events (events of LSV swelling) were identified and tagged. Each tagged fusion event was verified by observing the LSV across the imaging planes in order to exclude vesicles that move. Tagging was done exclusively in the middle slice of time-lapse stacks. In Glue-GFP and LifeAct-Ruby expressing SGs this was performed by viewing the Glue-GFP channel only, so as to minimize fusion detection bias. To allow sufficient imaging time of an LSV after a fusion event, so that the mode of exocytosis can be determined, the fusion events in the last 20 frames of the time-lapse are not tagged. At least 40 fusion events are tagged in a certain SG. In SGs that are secreting more rapidly (SGs with higher frequency of FC events for example), 2-3 cells (instead of the whole field of view) were selected at random and all the fusion events in these cells were tagged. Next – each fusion event was classified as “Crumpling”, “FC”, “KAR w/o Actomyosin”, “KAR w Actomyosin” or “Stalling”. Scenarios where the LSV does not lose volume after fusion until actin assembles (roughly within 30-60 seconds) and then content release proceeds normally were classified as crumpling events. Scenarios where content release occurs before actin was observed on the vesicle were classified as Full collapse events. Scenarios where there was fusion without any actin assembled on the LSV, or where content release did not occur were classified as KAR w/o Actomyosin. Scenarios where there was actin on the vesicle, but squeezing results in vesicle escape is classified as KAR w Actomyosin. Finally, scenarios where there was actin assembly followed by disassembly or where the LifeAct-Ruby intensified but without content release were classified as stalling. All events were identified in 2D, and verified by viewing the same object in other slices. Every quantification includes at least 120 fusion events from three or more different organisms. The time-lapse data sets used as biological repeats were not necessarily taken on the same day. Growth, sample preparation and imaging conditions were maintained between imaging days. GraphPad Prism version 8.4.2 (GraphPad software) was used for statistical analysis. We used multiple two tailed t-tests comparing each of the modes in the control to the same mode in the test groups.

### Sample preparation for FIB-SEM and correlative microscopy

Dissected *Drosophila* SGs were fixed using high-pressure freezing (Leica EM-ICE; Leica Microsystems). SG were placed in aluminum planchettes (Wolhwend; 0.3mm/0mm and 0.15mm/0.15mm, #1314 and # 1315 respectively) filled with Schneider’s *Drosophila* Medium supplemented with 10% BSA and 10% FBS to serve as cryoprotectant. Automatic freeze substitution (AFS) and resin embedding were performed in an EM-AFS2 mounted with EM-FSP (Leica Microsystems). HPF-fixed SGs were placed in 0.1% uranyl acetate in dry acetone at –90°C for 45h before the temperature was increased gradually, 2°C/h until it reached –45°C and remained there for an additional 40h, followed by 3 washes with acetone. Embedding with Lowicryl HM20 (EMS, cat#14340) was performed using gradual increase in resin concentration, 10%, 25%, 50% and 75%, 12h each. 10% and 25% infiltration were performed at –45°C, while 50% infiltration was performed after the temperature was increased to –35°C (0.8°C/h) and kept at –35°C for 75% infiltration step. The temperature was then increased to –25°C during the first 12h of 100% infiltration (0.8°C/h) and kept at –25°C for two additional rounds of exchange every 12-15h. The resin was then polymerized under UV for a total of 106h, while the temperature was increased to 20°C over the course of 10h (4.5°C/h). Blocks were left to for curing covered in foil until they turned from pink to transparent.

The blocks containing the SGs were trimmed using a razor blade from all sided, leaving a reduced block surface around the tissue. The block surface was sectioned using a diamond knife (35° ultra; Diatome) in an ultramicrotome (EM-UC7; Leica Microsystems), until the lumen of the gland was exposed and was visible on toluidine-blue stained sections. To help with targeting a region of interest (ROI), a mesh-like pattern was made manually on the polished surface with a fine razor (EMS; cat#72000).

### In-block FM for correlative imaging

The trimmed blocks were mounted in a drop of PBS in the center of a glass-bottom imaging plate (Cellvis; D35-10-1.5N) and fixed to position with plasticine. Blocks were imaged with an Olympus spinning disc confocal IX83 microscope (details under ‘**Confocal image acquisition’**). Iterative imaging and polishing were performed in cases where the regions of interest were more than ∼8μm deep from the polished surface.

### FIB-SEM sample preparation and imaging

Prior to FIB-SEM tomography, each block was mounted on an SEM stub (EMS; cat#75220) with a double-sided conductive carbon tape (EMS; cat#77816). The block edges were covered with additional strips of carbon tape and 2-4 strips of copper tape (EMS; cat#77802) and coated with a layer of colloidal silver liquid (EMS; cat#151105). Right before FIB-SEM imaging, the samples were sputter coated with an 8-10 nm layer of Ir using a high-vacuum compact coating unit (Safematic; CCU-010 HV).

Samples were mounted on a Crossbeam 550 FIB-SEM system (Carl Zeiss) and ROIs were located under 2kV, and either 5kV, 10kV or 15kV voltages, at 350pA or 1000pA before tilting the stage to 54° and adjustment of working distance to 5 mm. A 1μm-thick layer of Pt was deposited on top of the ROI using the ion beam (30kV, 0.3-1.5nA) and a trench was made (30kV, 3-30nA) with different dose factors (5-10), depending on the specific sample. The cross-section was then polished (30kV, 1.5-15nA) with the same dose factor before SEM imaging parameters were adjusted. Serial surface imaging was performed (30kV, 0.7-1.5nA) with the same dose factor as before, and SEM micrographs were acquired using either 2kV,0.35nA using a mixed signal from secondary electrons type 2 (SE2) and energy selective backscattered electrons (EsB) detectors with variable mixing ratios, depending on the sample or using 1.5kV, 1nA with EsB detection only. Scan speed and signal to noise increase were different depending on the specific block and volume acquired.

### FIB SEM and correlative image processing and analysis

The collected data was processed using Fiji (Schindelin et al., 2012). Stack alignment was performed using the registration plugin ‘Linear stack alignment with SIFT’, allowing translation only and without interpolation. Images were filtered with unsharp mask or local contrast enhancement (CLAHE) followed by smoothing. Fusion pore of vesicles before crumpling and after crumpling were measured in the plane where the pore is the widest, but in the narrowest point of the neck. Pore diameter measurements from FIB-SEM data (Fig. 1 E) includes 21 pores from 4 SGs for pre-crumpled LSVs and 22 pores from 3 SGs for crumpled LSVs, since the data is not evenly distributed, each pore was considered as an individual measurement.

For correlative microscopy, image stacks were binned in z between 2 to 4-fold, depending on the size of the stack, before loading to ICY (Chaumont et al., 2012) and Amira v2021.3 (Thermofisher Scientific) alongside the FM images of the block. Both datasets were roughly aligned in Amira as volumes to allow better identification of fiducials. Pairs of fiducials were identified in Amira volumes and placed on the data using eC-clem version 2 (in beta testing mode) (Paul-Gilloteaux et al., 2017). Affine transformation was applied on the FM stack based on 15-25 fiducial pairs and a complete error prediction of each point in the dataset was computed in the same way as described in (Scher et al., 2021; Potier et al., 2021). Last, to validate that the transformation did not drastically deform the FM information, the transformed FM stack was resliced to show xz planes, so as to observe the shape of the objects compared to the original FM stack.

### Statistics and reproducibility

Animal growing conditions, sample preparation, treatments and imaging conditions were kept similar between experiments. Modes of exocytosis characterization (frequencies of MC, FC, KAR and stalling events; Fig 2 C, 3 A, 4 B, 4 D, 6 B, 7 B and S5 B) are based on confocal live tissue imaging, *ex-vivo* of at least 3 SGs in which at least 40 fusion events per gland were followed. To characterize the changes in pore diameter, we used SRRF imaging to better visualize the pores of representative events. Where phenotypes were not apparent from the time-lapse sequences, at least n=9 events were quantified (as for MC and stalling in Fig. 1 D, 2 F and 3 C). Pore diameter measurements from FIB-SEM data (Fig. 1 E), were pooled from 4 individual data sets, each from a different gland (n=21 in pre-crumpled LSVs and n=22 in crumpled LSVs). Qualitative phenotypic characterization of knock-down experiments (summarized in Fig. 4 A) was based on at least 4 independent data sets using at least 2 different RNAi constructs for each gene (complete list of RNAi lines used in supplementary Table S1). GraphPad Prism version 8.4.2 (GraphPad software) was used for statistical analysis and for generating all plots. P values: ns≥0.05, *≤0.05, **≤0.01 ***≤0.001, ****≤0.0001. Further information on sample size, exact P values and the statistical tests used are mentioned in the figure legend and in the text for each experiment.

## Acknowledgments

We thank Prof. Sven Bogdan for the UAS-CIP4-EGFP *Drosophila* line, Dr. Helen Zenner for the MIM^Null^ *Drosophila* line and Prof Julie Brill for the Sgs3-DsRed *Drosophila* line. We thank the EM Unit of the Weizmann Institute for assistance. We thank all members of the O.A. and B.-Z.S. labs for fruitful comments and discussions. This research was supported by Israel Science Foundation (grant no. 706/20) to B.-Z.S., O.A., and E.D.S. and the Minerva Foundation with funding from the Federal German Ministry for Education and Research. O.A. also acknowledges funding from the Henry Chanoch Krenter Institute for Biomedical Imaging and Genomics, the Schwartz Reisman Collaborative Science Program, the Yeda-Sela Center for Basic Research, and the European Research Council (ERC) under the European Union’s Horizon 2020 research and innovation program (grant agreement no 851080). O.A. is an incumbent of the Miriam Berman presidential development chair. B.-Z.S. is an incumbent of the Hilda and Cecil Lewis Professorial Chair in Molecular Genetics.

## Author contributions

O.A., B-Z.S., T.B., and E.D.S. conceived and designed the experiments. T.B. carried out most of the experiments and analyzed the data. N.S. with help of Y.E.A performed and analyzed the FIB-SEM experiments. S.C. generated fly lines. O.A., T.B. and B-Z.S. wrote the manuscript with meaningful comments from all the authors.

## Supplementary figure legends

**Fig. S1.**
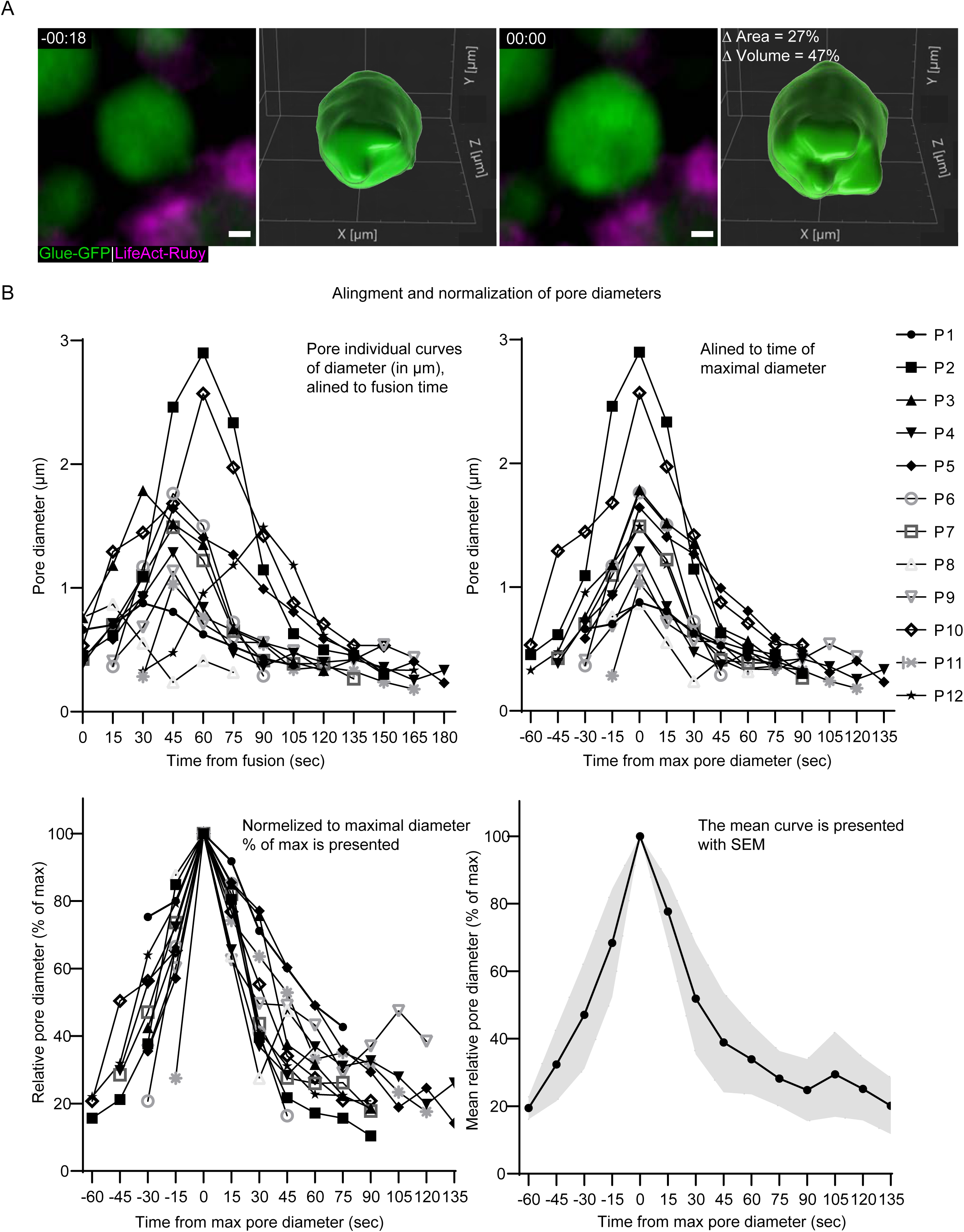
| LSV swelling, and pore diameter quantification and normalization. (A) 3D segmentation of an LSV before and after fusion showing vesicle swelling. Representative images (confocal Z-projection) and 3D segmentation of an LSV before (left) and after (right) fusion. The vesicle expands by 27% in 2D area and 47% in calculated 3D volume, implying that the observed increase in 2D is a result of LSV swelling and not LSV movement in or out of the imaging plane. Glue-GFP (Green) and LifeAct-Ruby (Magenta; UAS-based expression driven by c135-GAL4). Time mm:ss; relative to fusion. Scale bar is 1µm. (B) The step-by-step alignment and normalization plots used to produce the pore dynamics plot in **Fig. 1** **D**. (Top left) Pore diameter measurements (µm) are plotted over time (Seconds; from fusion). (Top Right) The curves are aligned over time (Seconds; relative to maximal pore diameter). (Bottom left) The curves are normalized (% of max pore diameter). (Bottom Right) The mean relative curve is presented with SEM (Gray). n=12.

**Fig. 2S.**
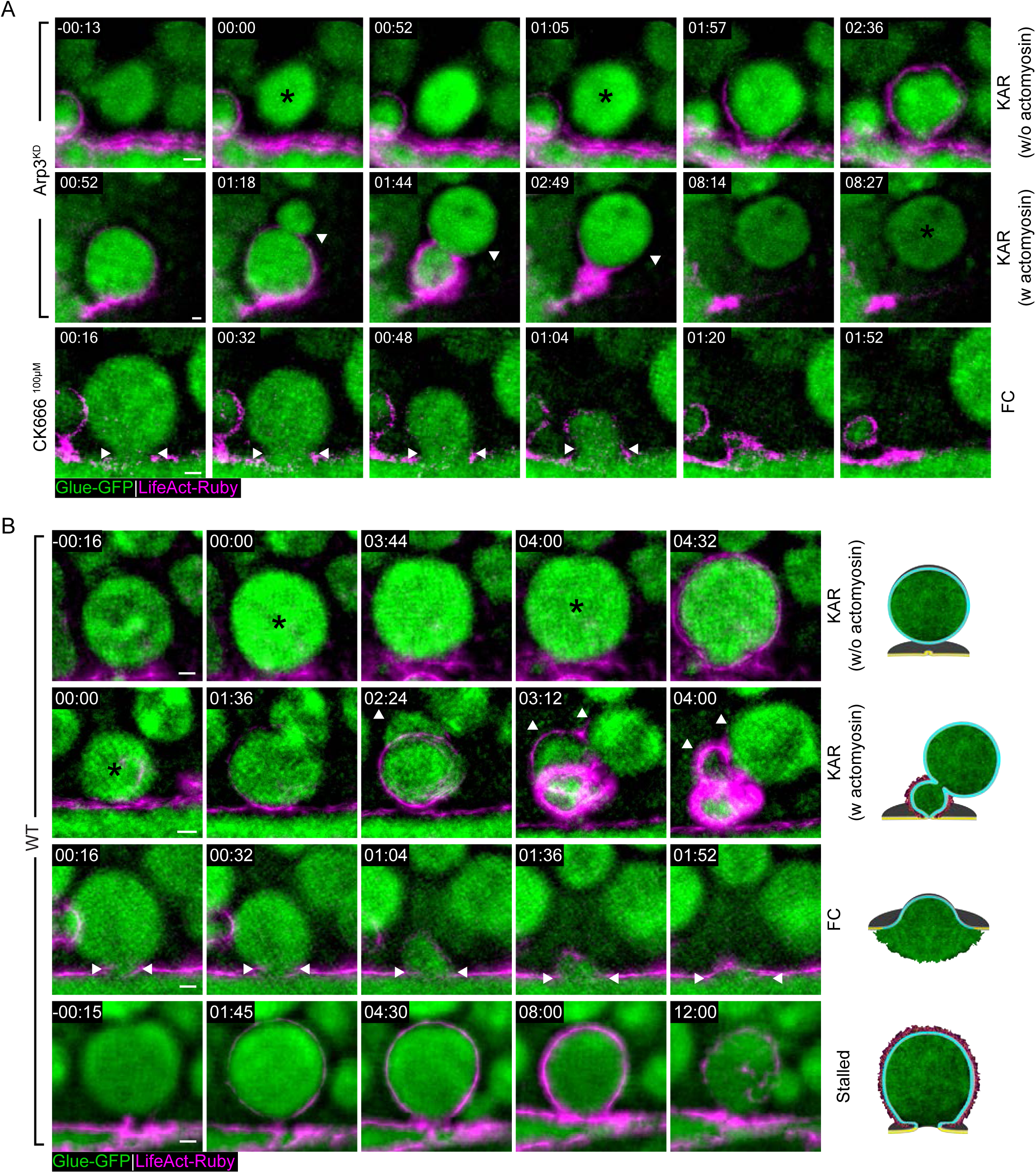
| Branched actin polymerization is essential for pore expansion and stabilization, and modes of exocytosis in the SG. (A) Time-lapse sequence of representative LSVs from Arp3^KD^ and CK666^100µM^ SGs undergoing KAR and FC (SRRF Z-projection), complementary to **Fig. 2** **A**, showing KAR in Arp3^KD^ SGs (top two rows) and FC in CK666^100µM^ treated SGs (bottom row). Glue-GFP (Green) and LifeAct-Ruby (Magenta). (B) Time-lapse sequence (SRRF Z-projection) of representative LSVs from WT SGs undergoing KAR (top two rows), FC (third row) and Stalling (bottom row). Fusion (Black asterisk), ‘jumping LSV’ (White arrow heads; in KAR w Actomyosin), fusion pore (dual white arrow heads; in FC). Glue-GFP (Green) and LifeAct-Ruby (Magenta). UAS-based expression in (A) driven by *fkh*-GAL4, in (B) driven by c135-GAL4. Time mm:ss; relative to fusion. Scale bars1µm.

**Fig. 3S.**
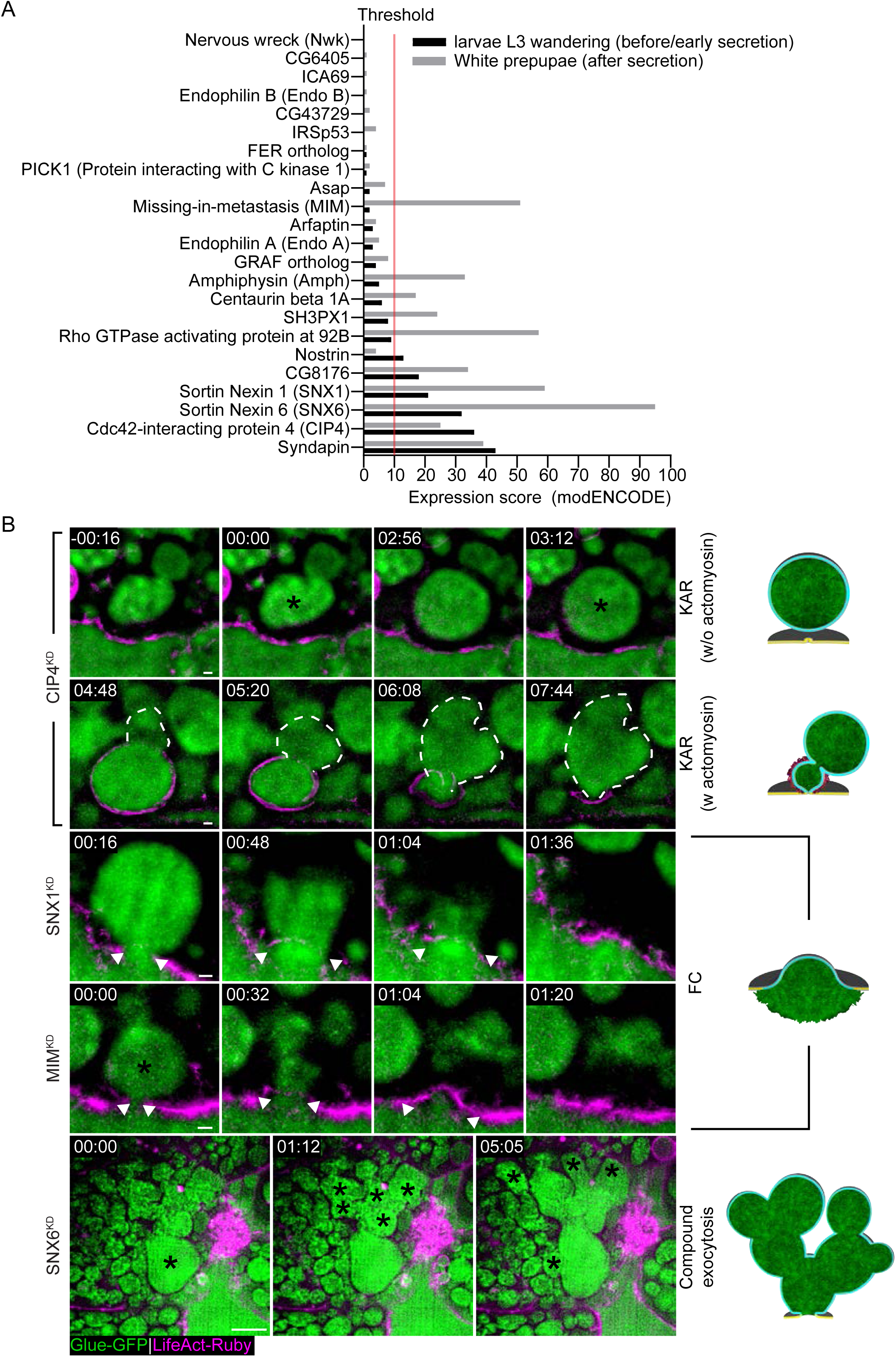
| Bar domain genetic screen and phenotypes. (A) *Drosophila* BAR domain gene expression scores (modENCODE, flybase), in SGs before (Black bars) and after the secretion phase (Gray bars). The 11 genes expressed above threshold (Red line), were included in our screen. (B) Time-lapse sequence of representative LSVs from BAR domain KD SGs undergoing KAR, FC and compound exocytosis (SRRF Z-projection). Fusion (Black asterisks), ‘jumping’ LSV (Dashed white line; in KAR w Actomyosin). Fusion pore (White arrow heads; in FC). In compound exocytosis we see that after fusion, LSVs fuse consecutively to an LSV fused to the cell membrane, creating a large ‘blob’. Glue-GFP (Green) and LifeAct-Ruby (Magenta). RNAi expressed under UAS control. UAS expression driven by *fkh*-Gal4. Time mm:ss; relative to fusion. Scale bars in KAR and FC 1µm, in Compound exocytosis 10µm.

**Fig. S4.**
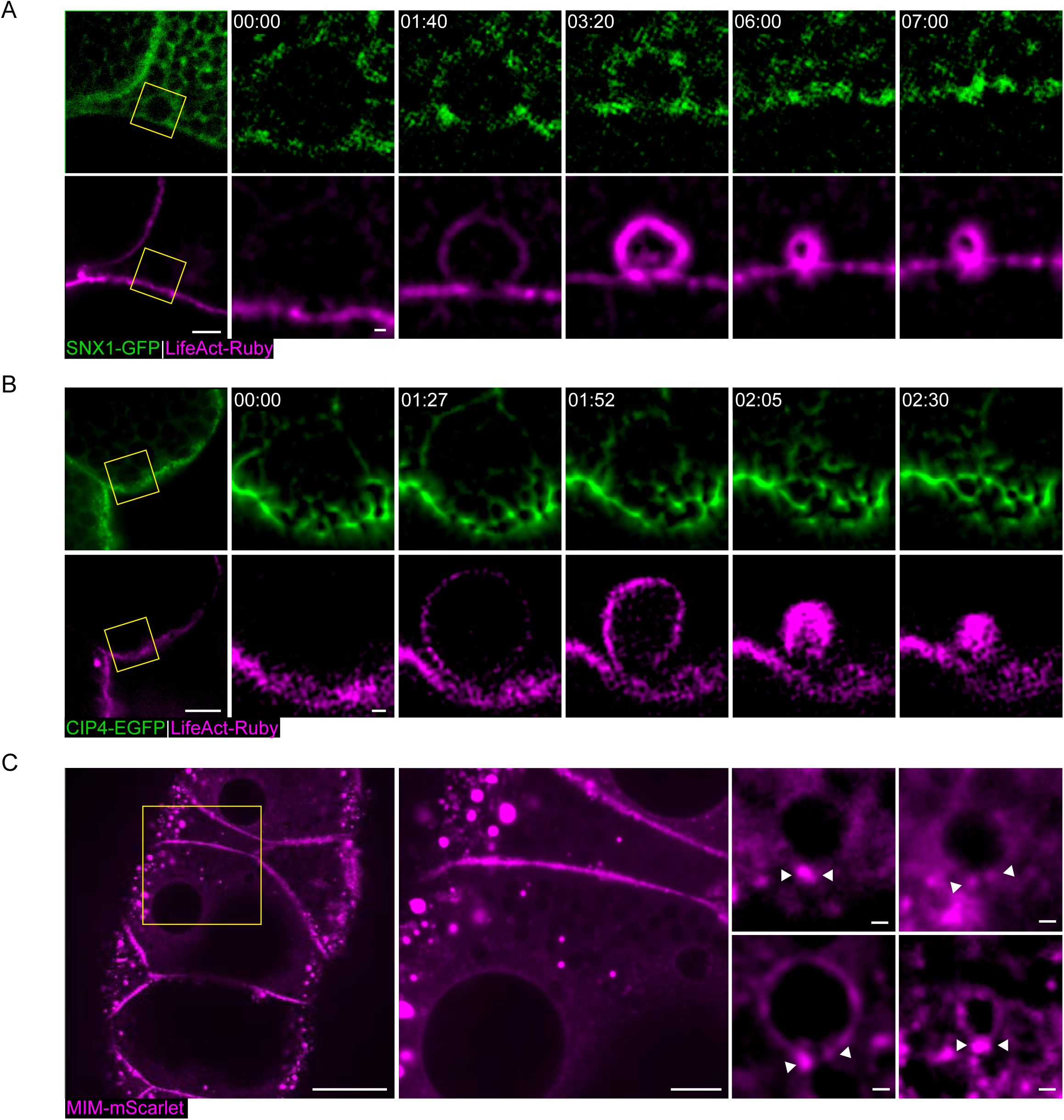
| SNX1-GFP, CIP4-EGFP and MIM-mScarlet localization. (A) Confocal Z projection and time-lapse sequence of a representative SG expressing SNX1-GFP (Green) and LifeAcrt-Ruby (Magenta), at increasing magnification. Yellow squares mark the magnified area. (Left) Representative image of a cell (confocal Z projection); SNX1 is seen on cell membranes and in the cytoplasm. (Right) Time-lapse sequence of representative secreting LSV (SRRF Z projection). Specific localization of SNX1-GFP to the fusion pore or to the LSV are not observed. (B) Confocal Z projection and time-lapse sequence of a representative SG expressing CIP4-EGFP (Green) and LifeAct-Ruby (Magenta), at increasing magnification. Yellow squares mark the magnified area. (Left) Representative image of a cell (confocal Z projection); CIP4-EGFP is seen mostly on the apical and lateral membranes of the cell. (Right) Time lapse of representative secreting LSV (SRRF Z projection). CIP4-EGFP is mostly apical and localizes to the LSV membrane after fusion, but specific localization to the pore was not detected. (C) Representative SG expressing MIM-mScarlet (**Fig. 6** **A;** Magenta). (Left) Overview of a large area in the gland (confocal Z projection). The yellow square shows the area enlarged in the middle image. (Middle) enlarged cell overview, Like MIM-Emerald (**Fig. 5** **A**) MIM-mScarlet localizes to cytoplasmic clusters and is absent from the apical membrane. Additionally, MIM-mScarlet localizes in the cytoplasm and to lateral membranes. (Right) Representative LSVs from different vesicles, in various stages of secretion (confocal Z projections). MIM-mScarlet is observed at and around the pore (white arrow heads) and is localized to LSV membrane during secretion. UAS-based expression in (A) – (C) driven by c135-GAL4. Scale bars in (A, Left) and (B, Left) 10µm, in (A, Right) and (B, Right) 1µm, in (C, Left) 40µm, in (C, Middle) 10µm, in (C, Right) 1µm.

**Fig. S5.**
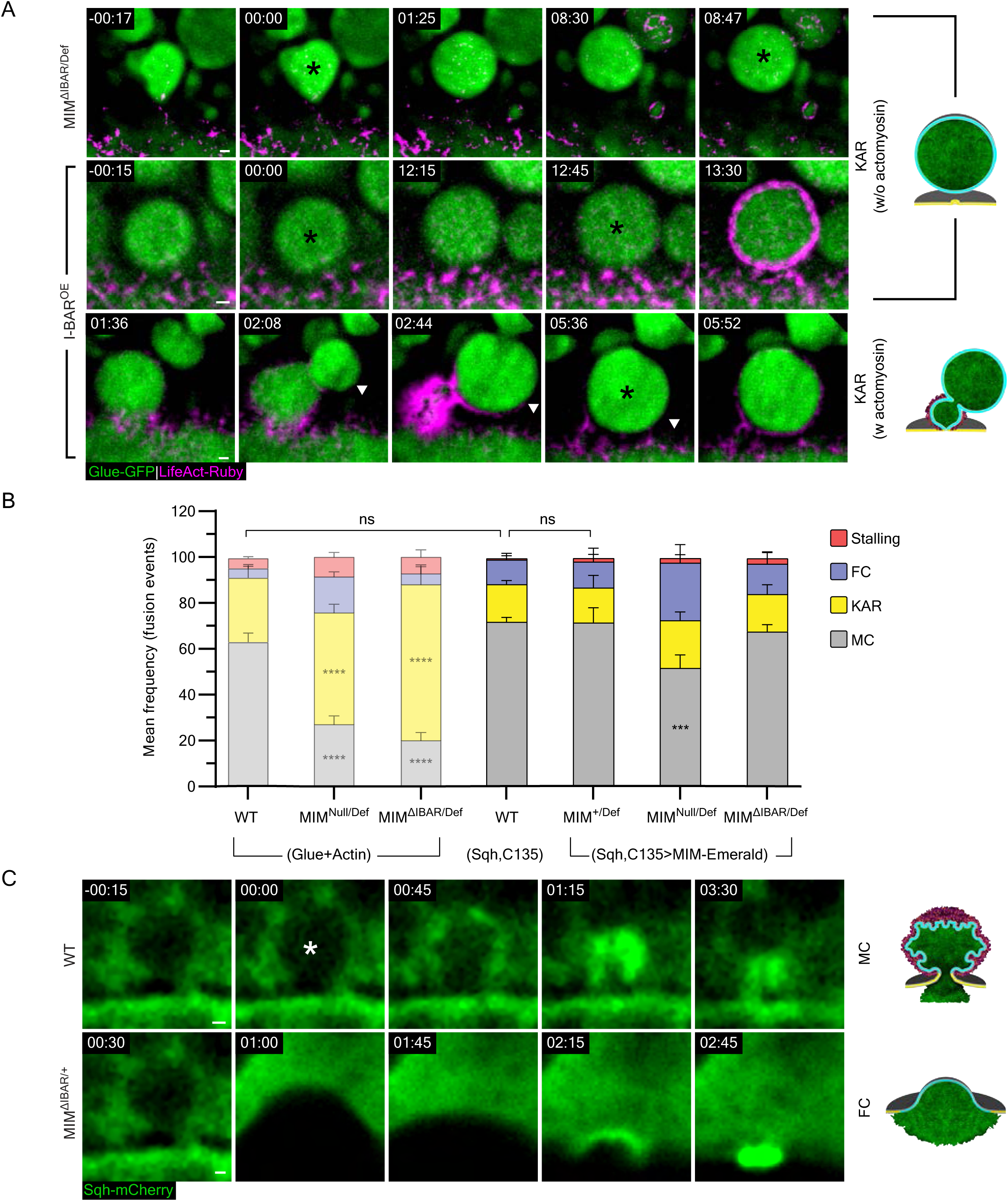
| Complementary examples to Fig. 7C, MIM-Emerald rescue experiment and myosin localization in MIM^ΔIBAR/+^. (A) Time-lapse sequence of representative LSVs from MIM^ΔIBAR/Def^ and I-BAR^OE^ SGs undergoing KAR w/o and w actomyosin (SRRF Z-projection). Complementary to **Fig. 6** **C**. Fusion (Black asterisk). Glue-GFP (Green) and LifeAct-Ruby (Magenta). (B) Mean frequency (%) of stalling, FC, KAR and MC in WT, MIM^Null/Def^ and MIM^ΔIBAR/Def^ using the Glue-GFP and LifeAct-Ruby markers and in MIM^+/Def^, MIM^Null/Def^ and MIM^ΔIBAR/Def^ expressing MIM-Emerald compared to WT glands using the Sqh-mCherry marker from **Fig. 6** **A and 7 A** (Semi-transparent bars). The majority of fusion events observed in WT SGs with Glue-GFP and LifeAct-Ruby markers result in MC events, MIM^Null/Def^ and MIM^ΔIBAR/Def^ SGs present the MIM loss-of-function phenotype and a high frequency of KAR events. WT SGs using the Sqh-mCherry marker do not significantly differ from the WT SGs using the Glue-GFP and LifeAct-Ruby markers. In MIM^+/Def^ SGs that also expressed MIM-Emerald the mode of exocytosis distribution did not vary significantly from the WT SGs using the Sqh-mCherry assay. When MIM-Emerald is expressed in MIM^Null/Def^ or MIM^ΔIBAR/Def^ SGs the loss-of-function phenotype is rescued and MC exocytosis is the major exocytosis mode observed. N(SGs)≥3, n(events)≥150. P values for MIM^Null/Def^: ***(MC)=0.009667 (two-tailed un-paired multiple t tests corrected using the Holm-Sidak method). (C) Time-lapse sequence of representative LSVs from WT SGs undergoing MC and MIM^ΔIBAR/+^ SGs undergoing FC (Confocal Z-projection). Fusion (White asterisks). Sqh-mCherry (Green). Time in (A) and (C) mm:ss; relative to fusion. UAS-based expression in I-BAR^OE^ (A) driven by c135-GAL4, in MIM^ΔIABR/Def^ (A) driven by *fkh*-GAL4, in (B) for the Glue-GFP and LifeAct-Ruby expressing SGs driven by *fkh*-GAL4 and for Sqh-mCherry expressing SGs driven by c135-GAL4. Scale bars in (A) and (C) 1µm.

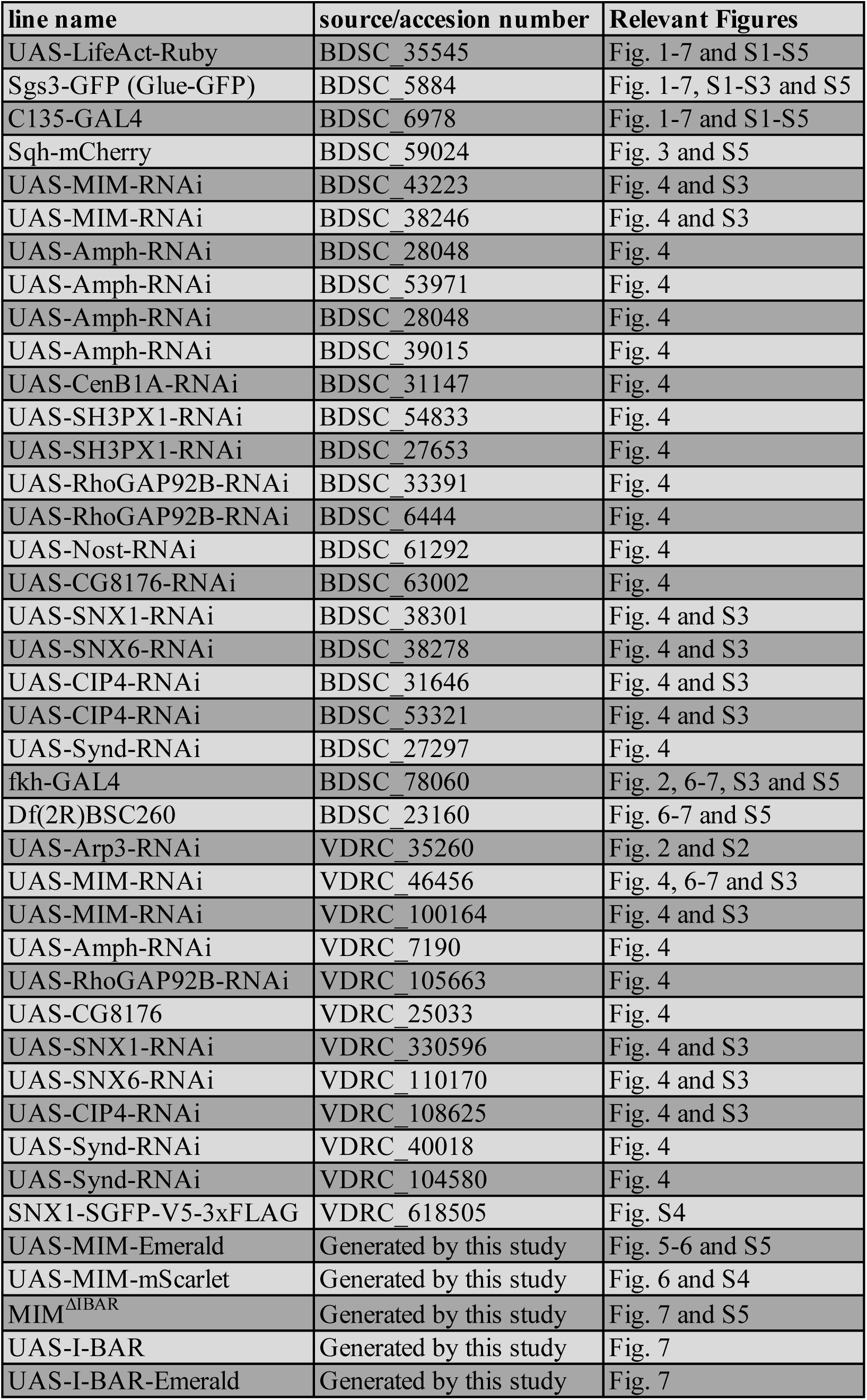

